# APOE4/4 is linked to damaging lipid droplets in Alzheimer’s microglia

**DOI:** 10.1101/2023.07.21.549930

**Authors:** Michael S. Haney, Róbert Pálovics, Christy Nicole Munson, Chris Long, Patrik Johansson, Oscar Yip, Wentao Dong, Eshaan Rawat, Elizabeth West, Johannes CM Schlachetzki, Andy Tsai, Ian Hunter Guldner, Bhawika S. Lamichhane, Amanda Smith, Nicholas Schaum, Kruti Calcuttawala, Andrew Shin, Yung-Hua Wang, Chengzhong Wang, Nicole Koutsodendris, Geidy E Serrano, Thomas G Beach, Eric M Reiman, Christopher K Glass, Monther Abu-Remaileh, Annika Enejder, Yadong Huang, Tony Wyss-Coray

## Abstract

Several genetic risk factors for Alzheimer’s Disease (AD) implicate genes involved in lipid metabolism and many of these lipid genes are highly expressed in glial cells. However, the relationship between lipid metabolism in glia and AD pathology remains poorly understood. Through single-nucleus RNA-sequencing of AD brain tissue, we have identified a microglial state defined by the expression of the lipid droplet (LD) associated enzyme *ACSL1* with ACSL1-positive microglia most abundant in AD patients with the *APOE4/4* genotype. In human iPSC-derived microglia (iMG) fibrillar Aβ (fAβ) induces *ACSL1* expression, triglyceride synthesis, and LD accumulation in an APOE-dependent manner. Additionally, conditioned media from LD-containing microglia leads to Tau phosphorylation and neurotoxicity in an APOE-dependent manner. Our findings suggest a link between genetic risk factors for AD with microglial LD accumulation and neurotoxic microglial-derived factors, potentially providing novel therapeutic strategies for AD.

---

Dr. Alois Alzheimer’s original description of what would later be known as Alzheimer’s disease (AD) included the identification of “many glial cells show[ing] adipose saccules” in the brains of patients with dementia^1^. The description of this glial-lipid pathological hallmark of AD was made alongside the descriptions of the plaque and tangle pathology commonly associated with AD, yet the glial-lipid hallmark of the disease has received relatively little attention in AD research. A recent meta-analysis of all genetic risk factors for AD identified through GWAS studies discovered genes involved in lipid processing and innate immunity as a statistically enriched category of genetic risk factors for AD, alongside the more characteristic categories of genes in amyloid and tau processing^2^, yet the role lipids and innate immunity play in AD risk remains poorly understood. APOE is one such lipid-related AD risk gene that is highly upregulated in human microglia in AD^3^ and human iPSC-derived microglia with APOE risk variants have more LDs.^4^. Aged mouse microglia accumulate LDs and exhibit a dysfunctional microglial state termed lipid droplet accumulating microglia (LDAM)^5^, and LDAM were also observed in chimeric human-mouse AD models^6^. LDs form in myeloid cells through the upregulation of lipid synthesis enzymes triggered by the engagement of toll receptors by innate immune triggers, such as bacteria^7^. LDs themselves have anti-microbial properties and are an evolutionarily conserved form of innate immune defense in macrophages^8^. Cholesterol-rich lysosomes and lipid droplets in dysfunctional microglia have also been observed in the context of demyelination mouse models and human iPSC models^9–12^. It remains unclear whether the lipid accumulating glial state originally described by Dr. Alzheimer in human AD brain tissue is influenced by lipid AD risk variants (e.g. *APOE*), if lipid accumulating glia reported in AD are similar to recently identified LDAM, and if lipid accumulating glia play a benign, protective or damaging role in AD pathogenesis.

## AD microglia have novel lipid transcriptional state defined by *ACSL1*

To investigate the transcriptional state of post-mortem human AD brain tissue in relationship to the APOE genotype we performed single-nucleus RNA-sequencing (snRNA-seq) on fresh-frozen frontal cortex tissue from individuals diagnosed with AD with the *APOE4/4* genotype, individuals with AD and an *APOE3/3* genotype and age and sex-matched control individuals with the *APOE3/3* genotype (**Fig. 1a, Extended Data Table 1**). This yielded ∼100,000 single nucleus transcriptomes with all major cell types of the brain represented (**Fig. 1b, Extended Data Fig. 1a, Extended Data Fig. 1a-f, Extended Data Fig. 2 a-p**). Differential gene expression analysis between control and AD-*APOE4/4* microglia revealed that the most significantly differentially expressed gene is *ACSL1,* which encodes a lipid processing enzyme (**Fig 1c,d, Extended Data Table 2**). ACSL1 is a key enzyme in LD biogenesis and overexpression of *ACSL1* is sufficient to induce triglyceride (TG) specific LD formation in multiple cell types^13, 14^ (**Fig 1e**). *ACSL1* was upregulated specifically in microglia in AD brain tissue compared to controls, and to a greater extent in *APOE4/4* compared to *APOE3/3* AD microglia (**Fig. 1f**). Subclustering all the microglia from this study revealed that ACSL1 positive microglia constitute a distinct state from homeostatic and DAM microglia, defined by the co-expression of additional metabolic state regulators such as *NAMPT*, and *DPYD* (**Fig. 1g,h, Extended Data Fig. 3 a-j**). Due to the set of LD-related genes in the *ACSL1* positive microglia cluster we refer to these *ACSL1* positive cells as lipid droplet accumulating microglia (LDAM). *APOE*4/4 AD brain tissue has the greatest percentage of LDAM, followed by *APOE3/3* AD, and the least amount of the LDAM microglia state is found in the aged-matched control brain tissue (**Fig. 1i**). Immunofluorescence microscopy staining of human AD brain tissue confirmed these ACSL1 abundance differences observed by RNA-seq (**Fig 1j,k**).

**Figure 1.**
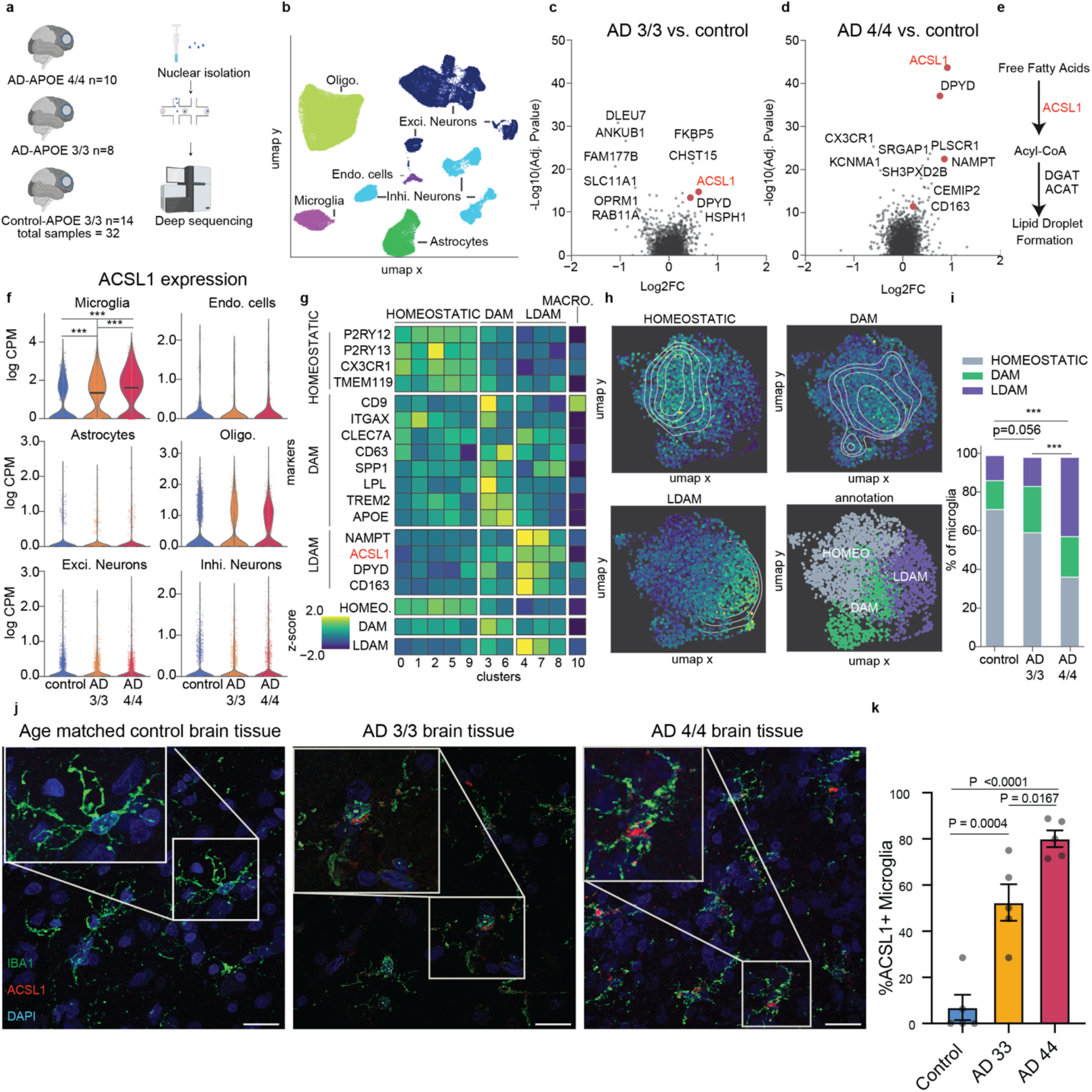
AD microglia have novel lipid transcriptional state defined by ACSL1. **a**, Schematic of single-nucleus RNA-seq cohort and workflow (see Methods). **b**, UMAP representation of all cells (n=100,317) from snRNA-seq, colored by annotated cell type. Data is shown after quality control and batch correction. **c,d,** Volcano plot representing MAST-based single-cell differential gene expression results (see Methods, Single-cell differential gene expression) of microglia from control subjects compared to microglia from subjects with AD and the APOE3/3 genotype (**c**) and from subjects with AD and the APOE4/4 genotype (**d**). Select lipid and metabolism-associated genes highlighted in red. **e**, Pathway diagram showing placement of differentially expressed gene *ACSL1* in pathway starting from free fatty acid to lipid droplet formation. **f**, Violin plots showing *ACSL1* expression across the cell types within the snRNA-seq dataset. Significance results indicate MAST-based adjusted p-values (see Methods, Single-cell differential gene expression). **g,** Normalized and z-scored gene expression levels of HOMEOSTATIC, DAM, and LDAM marker genes across the 11 subclusters identified within the microglia (**top**). HOMEOSTATIC, DAM, and LDAM signature scores are shown across the 11 identified subclusters. **h,** UMAP representation of microglia cells indicating the marker gene-based cell state annotation (**top left**), and the signature scores per cell for HOMEOSTATIC (**top right**), DAM (**bottom left**), and LDAM (**bottom right**) states. Contour lines indicate kernel density estimates of the signatures across the UMAP space. **i,** Bar plots indicating the percentage of cells from the three different cellular states (HOMEOSTATIC, DAM, and LDAM) across microglia from control, AD-APOE3/3, and AD-APOE4/4 groups. Chi-square test results indicate the significance of the percentage differences between the groups (*** indicates P<.0001). **j,** Representative immunofluorescence images of human frontal cortex adjacent to the tissue used in snRNA-seq experiments stained for microglia marker IBA1 (green), ACSL1 (red), and DAPI (blue) in an aged-matched healthy control subject (left), an AD APOE3/3 subject (middle), and an AD APOE4/4 subject. Scale bars (white, bottom right) 20 μm. **k,** Quantification of percentage of IBA1+ microglia positive for ACSL1. Each dot represents an average quantification for an individual subject. P-value determined by one-way ANOVA, error bar represents s.e.m.

**Figure 2.**
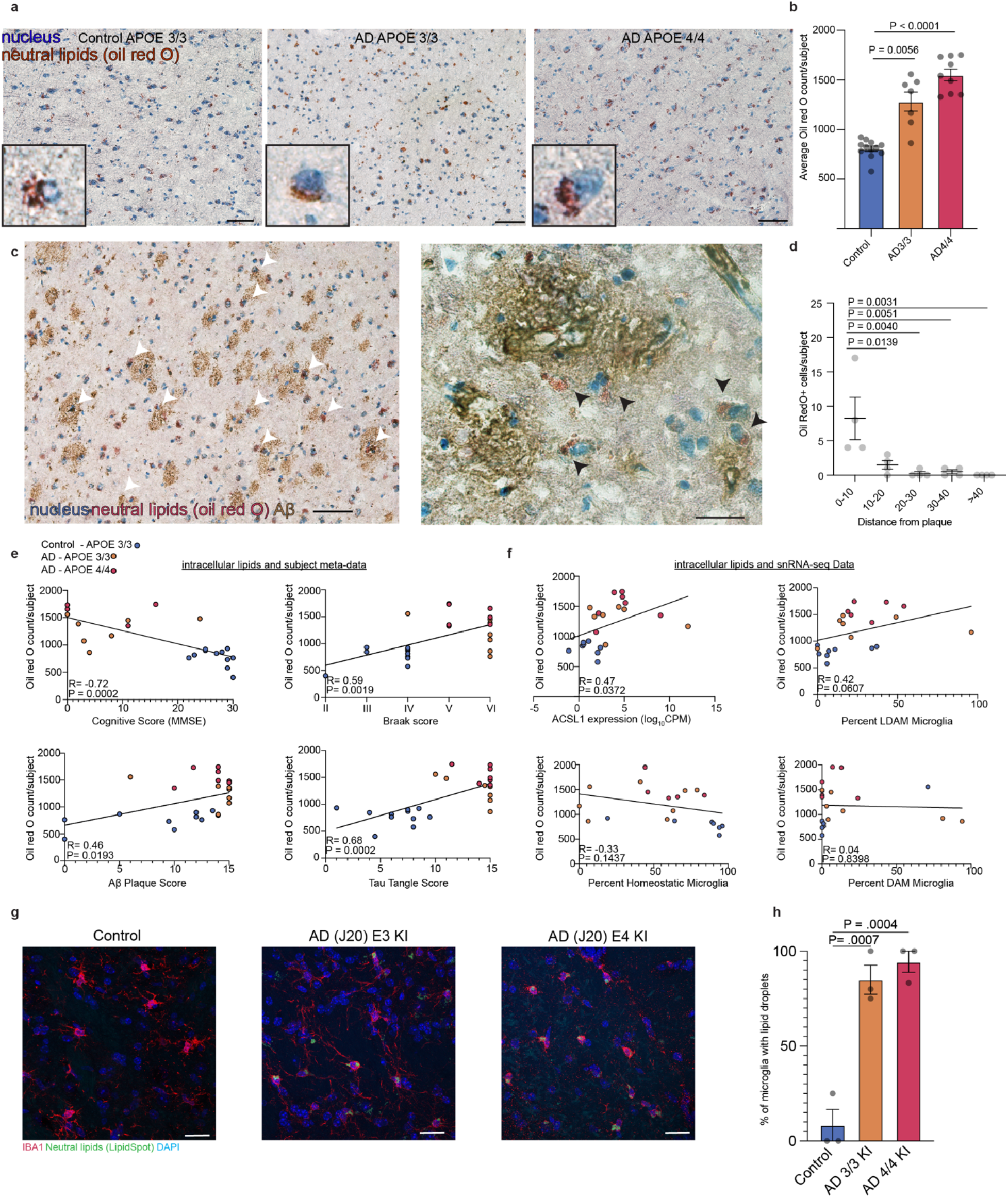
Intracellular lipid accumulation is linked to AD pathology. a, Representative Oil Red O staining image for control, AD *APOE3/3,* and AD *APOE4/4* human frontal cortex. Neutral lipids stained with Oil Red O (red) and nuclei stained with hematoxylin (blue). Scale bars (black, bottom right) 50 μm. b, Quantification of Oil Red O staining. Bar plots represent average Oil Red O counts per image for each subject category. Each dot represents average Oil Red O counts for an individual subject averaged over five 20x image fields per individual. P-value determined by one-way ANOVA, error bar represents s.e.m. c, Oil Red O staining of *APOE4/4* AD subjects with IHC staining for Aβ. White arrow heads represent Oil Red O positive cells in or around Aβ plaques. Scale bars (black, bottom right) 50 um (left). High magnification of representative Oil Red O stain with IHC staining for Aβ in AD *APOE4/4* subject. White arrowheads represent Oil Red O positive cells in or around Aβ plaques. Scale bars (black, bottom right) 20 μm (right). d, Quantification of the frequency of Oil Red O positive cells in various distances from Aβ plaques. Each dot represents an individual subject. P-value determined by one-way ANOVA, error bar represents s.e.m. e, Scatter plot of average Oil Red O counts per subject averaged over five 20x image fields per individual with individual subject’s meta-data. Subject category colored blue for control, orange for AD *APOE3/3* subjects, and red for AD *APOE4/4* subjects. P-values determined by spearmen correlation. f, Scatter plot of average Oil Red O counts per subject averaged over five 20x image fields per individual with individual subject’s snRNA-seq data. Subject category colored blue for control, orange for AD *APOE 3/3* subjects and red for AD *APOE4/4* subjects. P-values determined by Spearmen correlation. g, Representative immunofluorescence images of mouse hippocampus tissue stained for microglia marker IBA1 (red), neutral lipids (LipidSpot: green), and DAPI (blue) in control age-matched non-transgenic mice (left), AD mouse model (J20) with human APOE3 knock-in (middle), and AD mouse model (J20) with human *APOE4* knock-in (right). Scale bars (white, bottom right) 20 μm. h, Quantification of average percent of IBA1 positive microglia with neutral lipid dye (LipidSpot). Each dot represents individual biological replicate. P-value determined by one-way ANOVA, error bar represents s.e.m.

**Figure 3.**
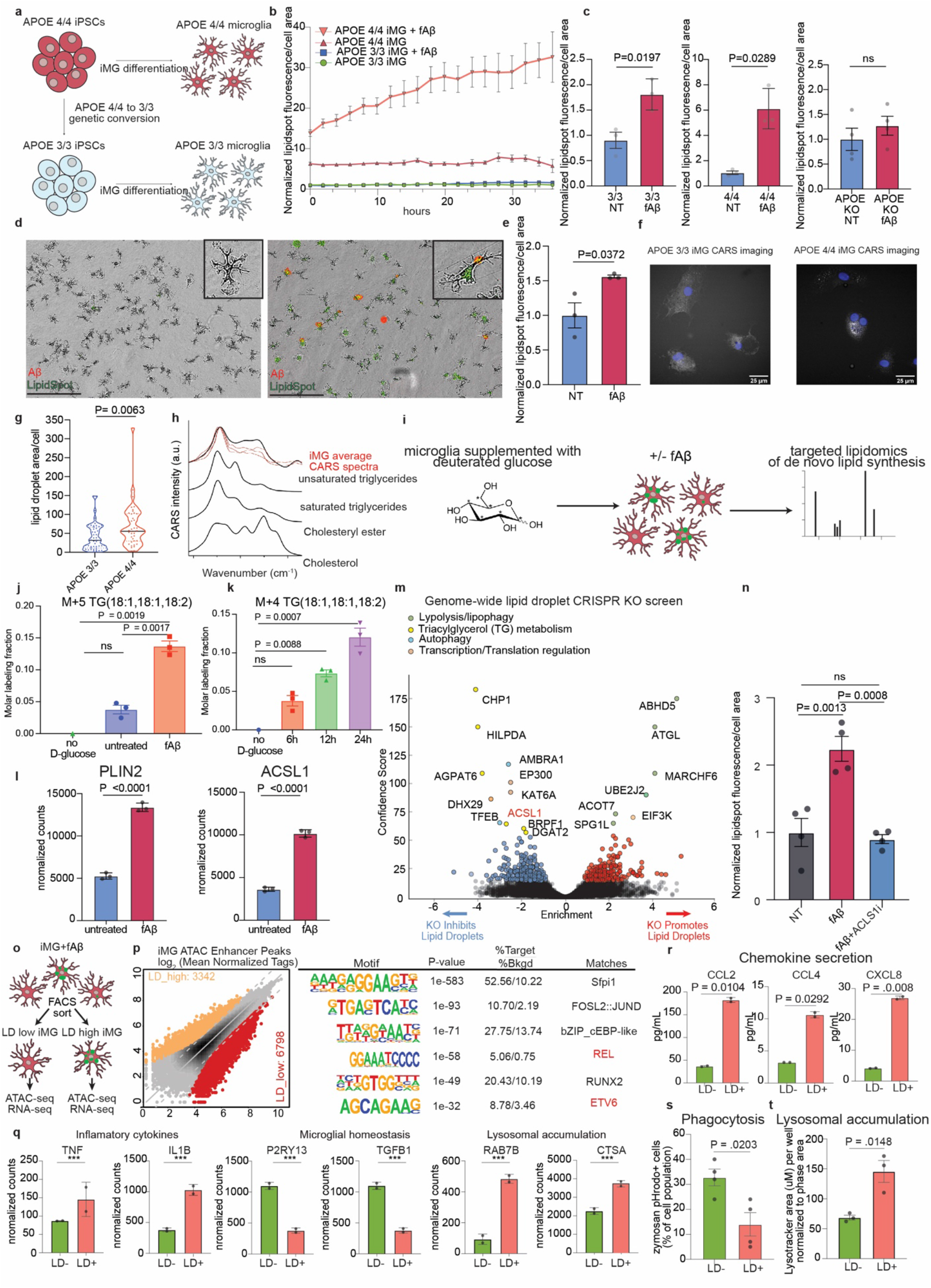
iMG upregulate ACSL1 and increase TG lipid synthesis upon fAβ challenge. **a**, Schematic of isogenic *APOE3/3* and *APOE4/4* iPSCs and differentiation into iPSC-derived microglia (iMG). **b**, Live cell imaging of untreated *APOE3/3* and *APOE4/4* iMG and fAβ treated *APOE3/3* and *APOE4/4* iMG stained with a green lipid fluorescent dye (Lipidspot). The y axis represents mean green fluorescence per cell normalized to untreated APOE3/3 iMG at the first time point and the x-axis represents imaging time points in hours (n=3 replicate wells per condition, error bars represent s.e.m.). **c**, Average Lipidspot green fluorescence per cell normalized to untreated of final timepoint in **b**. Individual dots represent replicate wells (n=3, error bars represent s.e.m, p-value calculated by unpaired, two-sided t-test). **d**, Primary rat microglia cultured under serum-free conditions untreated (left) or treated with fAβ (red) and stained with a green lipid fluorescent dye (Lipidspot). Scale bar 200 μm. **e**, Average Lipidspot green fluorescence per cell normalized to untreated images in **d**. Individual dots represent replicate wells (n=3, error bars represent s.e.m, p-value calculated by unpaired, two-sided, t-test). **f**, Representative CARS images of *APOE4/4* and *APOE3/3* iMG treated with fAβ, Scale bars (white, bottom right) 20 μm. **g**, Quantification of lipid droplet measurements from CARS microscopy of APO4/4 and APOE iMG treated with fAβ. Each dot represents lipid measurements from individual cell measurements. P-value calculated by unpaired t-test. **h**, Example CARS spectra from fAβ treated iMG (red) and reference spectra for common lipid species (black). **i**, Schematic of lipidomics measurement of C13-D-Glucose incorporation into TG over time in BV2 cells treated with 5uM fAβ. n=3, error bars represent s.e.m, p-value calculated by on-way ANOVA. **j**, Incorporation of C13-D-Glucose into triglycerides in microglia after fAβ treatment or untreated cells compared with cells without C13-D-Glucose as control. Each dot represents an individual replicate. n=3, error bars represent s.e.m, p-value calculated by one-way ANOVA. **k**, Incorporation of C13-D-Glucose into triglycerides in microglia after fAβ treatment over time compared with cells without C13-D-Glucose as control. Each dot represents an individual replicate. n=3, error bars represent s.e.m, p-value calculated by one-way ANOVA. **l**, Normalized gene expression counts for significant differentially expressed genes between untreated and fAβ treated APOE4/4 iMG (n= 3 replicate wells, error bars represent s.e.m, P-values determined by DEseq2. **m**, Volcano plot of Genome-wide CRISPR KO screen for BODIPY high and BODIPY low cells using the human monocyte U937 cell line. The confidence score and effect score are determined by CasTLE. Genes passing a 10% FDR cutoff are highlighted in red and blue. **n**, Average Lipidspot green fluorescence per cell normalized to untreated cells with 5uM fAβ, and 5uM fAβ with 1uM ACSL1 inhibitor (Triacin C). Individual dots represent replicate wells (n=4, error bars represent s.e.m, p-value calculated by unpaired, two-sided t-test). **o**, Schematic of ATAC-seq and RNA-seq experiments to characterize lipid droplet high and lipid droplet low iMGs. **p**, Genome-wide comparison of open ATAC-seq enhancer peaks comparing LD low and LD high iMG (left). *De novo* Motif analysis of differential peaks using HOMER. Motifs enriched in lipid-associated macrophages are highlighted in red. **q**, Normalized gene expression counts for significant differentially expressed genes between lipid droplet high from lipid droplet low APOE4/4 iMG (n= 3 replicate wells, error bars represent s.e.m, P-values determined by DEseq2, *** P < 0.0001). **r**, Measurement of secreted chemokines in cell culture media after FACS separation of lipid droplet positive from lipid droplet negative *APOE4/4* iMG following 18-hour Aβ treatment. Individual dots represent replicate wells (n=2, error bars represent s.e.m, p-value calculated by unpaired, two-sided t-test). **s**, Average percent of pHrodo zymosan red positive of total cell population with or without lipid droplets (Lipidspot) after 18-hour Aβ treatment. Individual dots represent replicate wells (n=3, error bars represent s.e.m, p-value calculated by unpaired, two-sided t-test). **t**, Average of percent of lysotracker red positive of total cell population with or without lipid droplets (Lipidspot) after 18-hour Aβ treatment. Individual dots represent replicate wells (n=3, error bars represent s.e.m, p-value calculated by unpaired, two-sided t-test).

## Intracellular lipid accumulation is linked to AD pathology

To measure intracellular lipid accumulation AD and control brain sections were stained with Oil Red O, a dye for neutral lipids. The *APOE4/4* AD patient brains showed an abundance of perinuclear Oil Red O positive lipid bodies that resemble LD and are similar to Alzheimer’s original description of “adipose saccules” in glial cells of post-mortem patient brain tissue (**Fig. 2a**). These lipid bodies are most prevalent in AD brain tissue with a slight but not significant increase in *APOE4/4* AD subjects compared to *APOE*3/3 AD subjects. (**Fig. 2b, Extended Data Fig. 4a**). Oil Red O positive cells are often found near to or at the core of amyloid-beta plaques (**Fig. 2c,d, Extended Data Fig. 4b,c**). In a similar fashion, ACSL1-positive microglia are often observed near amyloid-beta plaques (**Extended Data Fig. 4d**), suggesting that the cells containing lipid bodies near plaques could be ACSL1-positive microglia. However, the identification of the cell type(s) that accumulate these lipids necessitates simultaneous staining for lipids and multiple protein markers, which is challenging in aged human histological brain sections with current technology.

**Figure 4.**
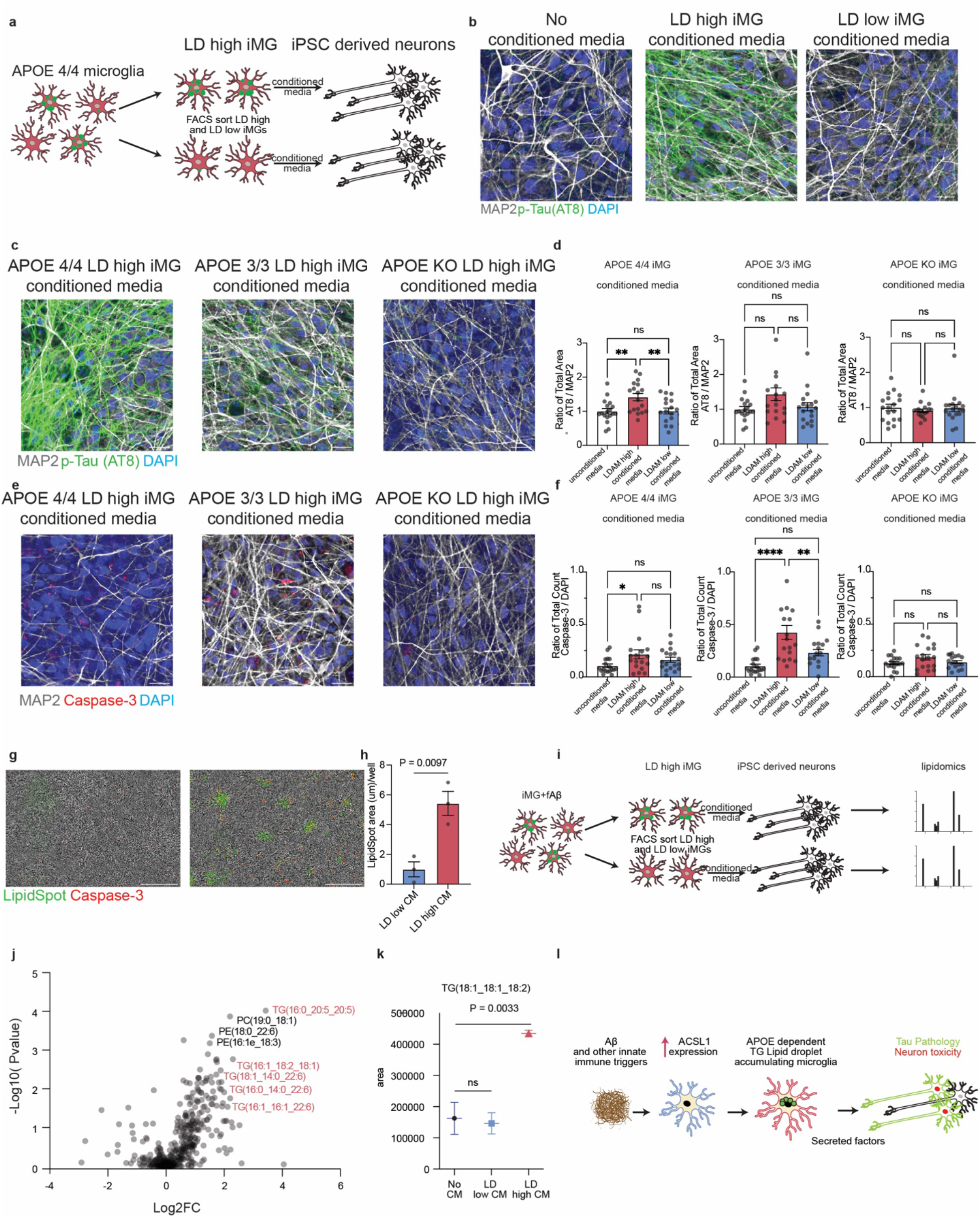
LD-containing microglia induce Tau phosphorylation and apoptosis in neurons. **a**, Schematic of LDAM-specific conditioned media exposure to neurons. **b**, Representative immunofluorescence images of iPSC-derived neurons exposed to no conditioned media (left), lipid droplet high APOE4/4 iMG conditioned media (middle), and lipid droplet low APOE 4/4 iMG conditioned media (left). Cells were stained for DAPI, MAP2 (grey), and phosphorylated Tau (AT8: green). Scale bars (white, bottom right) represent 20 μm. **c**, Representative immunofluorescence images of iPSC-derived neurons exposed to lipid droplet high APOE4/4 iMG conditioned media (left), lipid droplet high APOE3/3 iMG conditioned media (middle), lipid droplet high APOE KO iMG conditioned media (right). Cells were stained for DAPI, MAP2 (grey), and phosphorylated Tau (AT8: green). Scale bars (white, bottom right) represent 20 μm. **d**, Quantification of images as presented in **c** by measuring the AT8 fluorescence area normalized to the MAP2 florescence area. Each dot represents a random filed image (N=18) across 3 replicate wells, error bars represent s.e.m, p-value calculated by ANOVA. **e**, Representative immunofluorescence images of iPSC-derived neurons exposed to lipid droplet high *APOE4/4* iMG conditioned media (left), lipid droplet high *APOE3/3* iMG conditioned media (middle), lipid droplet high APOE KO iMG conditioned media (right). Cells were stained for DAPI (blue), MAP2 (grey), and cleaved Caspase-3 (red). **f**, Quantification of images as presented in **e** by counting cleaved caspase-3 fluorescence regions normalized to DAPI. Each dot represents a random filed image (N=18) across 3 replicate wells, error bars represent s.e.m, p-value calculated by ANOVA. **g**, Representative immunofluorescence images of neurons exposed to lipid droplet low *APOE4/4* iMG conditioned media (left) and lipid droplet high APOE4/4 iMG conditioned media (right). Cells were stained for neutral lipids (LipidSpot: green) and activated caspase-3 dye (red). **h**, Quantification of average LipidSpot green fluorescence area cell per well normalized to the lipid droplet low iMG conditioned media treated neurons. Individual dots represent replicate wells (n=3, error bars represent s.e.m, p-value calculated by unpaired, two-sided t-test). **i**, Schematic of lipidomics experimental design of neurons treated with conditioned media. **j**, Volcano plot representing lipids detected in neurons after treatment of lipid droplet high iMG conditioned media compared to lipid droplet low iMG conditioned media. TG species are highlighted in red. **k**, Representation of lipidomic measurements from one lipid species detected in lipidomic analysis. Individual dots represent replicate wells (n=3, error bars represent s.e.m, p-value calculated by one-way ANOVA. **l**, Schematic of the proposed role of lipid droplet accumulating microglia in neurodegeneration.

The number of lipid bodies is negatively correlated with cognitive performance, as measured by the mini-mental state exam (MMSE), and positively correlated with Aβ plaque levels and Tau pathology levels (**Fig. 2e**). Since Oil Red O staining and snRNA-seq were done on the same samples we were able to correlate gene expression with the abundance of lipid bodies for each sample. Reassuringly, the expression of *ACSL1* by microglia positively correlated with the relative numbers of lipid bodies (**Fig. 2f**). Mirroring these observations in the human AD tissue, there are more lipid droplet-positive microglia in the J20/*APOE3* and J20/*APOE4* models of AD^15^ compared to age-matched WT mice (**Fig. 2g,h**).

## iMG upregulate ACSL1 and undergo TG lipid synthesis upon fAβ challenge

To directly test whether the *APOE* genotype contributes to LDs accumulation in microglia, *APOE4/4* and isogenic *APOE3/3* iPSCs^16^ were differentiated into microglia (iMG) as previously described^17–19^ (**Fig. 3a**). Live cell microscopy of iMG with a fluorescent dye for neutral lipids (Lipidspot) showed greater LD accumulation in *APOE4/4* iMG compared with isogenic *APOE3/3* iMG, similar to recent reports in isogenic iPSC derived microglia^4^ and astrocytes^20^. However, treatment of iMG with fAβ led to a strong increase in LD accumulation that was exacerbated by the presence of the *APOE4* AD risk allele. The effect of fAβ on LD accumulation was absent in the *APOE* KO background (**Fig. 3b,c**). To ensure the LD induction in microglia by fAβ is not unique to these iPSCs lines or the differentiation protocol, primary rat microglia were treated with fAβ, and LD accumulation was also observed (**Fig. 3d,e**). The induction of LDs by fAβ was also observed in primary human macrophages and the mouse BV2 microglial cell line (**Extended data Fig. 5d**). In addition to lipid dyes to measure LD accumulation, transmission electron microscopy (TEM) indicated an increase in lipid droplet levels in iMG upon fAβ challenge (**Extended data Fig. g-h**). Coherent anti-Stokes Raman scattering (CARS) imaging was performed on iMG to confirm differential lipid accumulation between the *APOE* genotypes upon the fAβ challenge. Analysis of the CARS imaging of fAβ treated iMGs revealed that the lipid droplet spectra overlap with unsaturated triglyceride (TG) spectra (**Fig. 3f-h**). To investigate if these lipids are synthesized *de novo* in response to fAβ, BV2 microglia were grown with deuterated glucose (d-glucose) (**Fig 3i**). Lipidomic analysis showed a time-dependent increase in TG incorporation of d-glucose upon fAβ challenge (**Fig 3j,k**). In accordance with this accumulation of lipid droplets, the gene expression of LD-associated genes *PLIN2*^21^ and *ACSL1*^13, 14^ are upregulated upon fAβ challenge in iMGs (**Fig. 3l**) and to a greater extent in the *APOE4/4* background (**Extended data Fig. 3i,j**). A previously published dataset shows that *ACSL1* is highly upregulated by the innate immune trigger LPS in an independent human iMG chimeric mouse model^22^ (**Extended data Fig. 5k**), suggesting that *ACSL1* upregulation in microglia is the consequence of a broader response to innate immune triggers. To assess which specific lipid synthesis genes in the human genome play a role in lipid droplet accumulation, we performed a genome-wide CRISPR KO screen in the monocyte cell line U937 by FACS. This screen revealed regulators of TG metabolism as being a top category of genes required for LD accumulation and *ACSL1* as one of the most significant genes required for LD formation (**Fig. 3m, Extended data table 3**). An ACSL1 inhibitor (Triacin C) reversed the accumulation of LD in APEO4/4 iMG upon fAβ challenge (**Fig. 3n**).

To assess the transcriptomic and epigenetic state of microglia with LD accumulation we performed FACS sorting of LD high and LD low iMG followed by ATAC-seq and RNA-seq (**Fig 3o, Extended data table 4**). We detected 3,442 peaks that were gained in LD-high microglia compared to LD-Low microglia. ATAC-seq peaks at enhancer regions for LD-high were highly enriched for microglia lineage-determining transcription factors for PU.1. In addition, peaks specific for LD-high microglia showed enrichment for motifs related to the NF-κB family of transcription factors (e.g. REL, ETV6) **(Fig. 3p**). This epigenetic signature is also seen in macrophages in atherosclerosis models^23^ and adipose-associated macrophages^24^, but this signature is not seen in DAM microglia^25^. RNA-seq results indicated that LD-high microglia had higher expression of NF-κB associated proinflammatory cytokines (e.g *TNFA, IL1B*) and lower expression of microglial homeostasis markers compared to LD-low microglia (**Fig. 3q**). In accordance with these RNA-seq results, phenotypic measurements of LD-containing *APOE4/4* iMGs indicate that they are dysfunctional in phagocytosis, accumulate lysosomes, and secrete inflammation-associated chemokines as measured in the cell culture media. These results are in accordance with recent reports describing mouse LDAM to represent a dysfunctional and inflammatory microglial state^5^ (**Fig. 3 r-t**). Interestingly, among the top differentially-expressed genes between *APOE3/3* LDAM and *APOE4/4* LDAM, we identified the anti-bacterial LD-associated protein Cathelicidin or *CAMP* (**Extended data Fig. 4a, Extended data table 5**), which is found in LD with antimicrobial properties in macrophages exposed to bacteria^8^.

To discover genetic modifiers of lipid droplet accumulation in iMG by fibrillar Aβ, we performed a CRISPR KO screen in *APOE4/4* iMG with a library of ∼20,000 sgRNAs targeting the “druggable genome” of ∼2,000 genes representing all human kinases, phosphatases and known drug targets (**Extended Data Fig. 7a**). The top hit from this screen was *PIK3CA*, a catalytic subunit of PI3 kinase (**Extended Data Fig. 7b, Extended data table 6**). Interestingly, the second top hit was S100A1, an AD risk gene downstream of the LPS and TLR4 response in macrophages. PI3 kinase inhibition is a known modulator of lipid droplets in mouse macrophages^26^, but this has not been previously shown in human microglia. We tested if inhibition of PI3 kinase might reduce lipid droplet accumulation in iMGs. Indeed, the small molecule PI3K inhibitor GNE-317 dramatically reduced lipid droplet formation in APOE4/4 iMGs exposed to fAβ quantified by live microscopy and PLIN2 staining (**Extended Data Fig. 7c,d**). In addition, PI3K inhibition with GNE-317 reversed the lysosomal accumulation and inflammatory cytokine secretion observed in iMGs with high levels of lipid droplets (**Extended Data Fig. 7 e,f**). To further investigate the effects of GNE-317 we performed RNA-seq of APOE4/4 iMGs treated with fAβ in the presence or absence of the drug. GNE-317 reduced the expression of genes involved in lipid synthesis and reduced the genes indicative of a dysfunctional microglia state, such as inflammatory cytokine production and lysosomal accumulation (**Extended Data Fig. 7g,h**), and increased expression of genes involved in lipid degradation, microglial homeostasis, and neuroprotective growth factors, such *BDNF* and *FGF1* (**Extended Data Fig. 7 i,j, Extended data table 5**). Upon GNE-317 treatment, genes in the PI3K/mTOR and autophagy pathways exhibit changes in expression, and we also observed an increase in LC3B protein levels (**Extended Data Fig. 7d k-n**). This may indicate that GNE-317 treatment reduced lipid droplet levels through increasing autophagy, a mechanism that is known to regulate lipid droplet _levels_^27,28,29^.

## LD containing microglia induce tau phosphorylation and caspase activation in neurons

To investigate the effect of LDAM on neurons, *APOE4/4* iMG were FACS sorted into LD-high (top 10% BODIPY signal) and LD-low fractions (bottom 10% BODIPY signal) and cultured for 12 hours in neurobasal media to create conditioned media. *APOE*4/4 iPSC-derived human neurons were then grown in complete media containing 10% of the LD-high or LD-low *APOE*4/4 iMG conditioned media, as well as an untreated control condition (**Fig. 4a**). This approach takes inspiration from recent work showing the deleterious effects of conditioned media on neurons from astrocytes^30^ and microglia^4^. To investigate whether LDAM conditioned media induced hallmarks of AD pathology, the human iPSC-derived neurons were then stained with monoclonal antibody AT8 to detect phosphorylated Tau(pTau). Strikingly, only the LD-high iMG conditioned media induced high levels of pTau in the iPSC-derived neurons, whereas the LD-low iMG conditioned media induced similar levels of pTau in the iPSC-derived neurons as the untreated condition (**Fig. 4b**). This effect was similar when conditioned media from *APOE*3/3 and *APOE*4/4 iPSC-derived iMG were used to treat the human neurons, but absent when conditioned media from *APOE* KO iMGs were used (**Fig. 4 c,d**). Likewise, conditioned media from *APOE*3/3 and *APOE*4/4 iMGs with a higher number of LDs induced caspase activation in human neurons, while conditioned media from *APOE* KO iMG had no effect (**Fig 4. f,g**).

Intriguingly, human neurons treated with the LD-high conditioned media showed an increase in LipidSpot staining **(Fig. 4h)**. To investigate which lipids accumulate in iPSC-derived neurons upon LDAM conditioned media treatment, we performed lipidomics on neurons exposed to LD-high and LD-low conditioned media. The iPSC-derived neurons exposed to LD-high conditioned media contained higher levels of the TG lipid species that accumulate in iMG upon fAβ challenge (**Fig 4. j-k, Extended data table 7**). ^13^C-labeled TAG lipids synthesized in microglia were also observed in mouse neurons treated with conditioned media from BV2s grown in ^13^C-labeled glucose, however, this was not robustly detected across multiple replicates perhaps due to the low isotopic enrichment of these lipids (**Extended data Fig 8a**).

## Discussion

Here we report a novel microglial state present in human brains defined by the accumulation of lipids with concomitant upregulation of genes involved in lipid synthesis; these cells are phenotypically similar to previously reported mouse LDAM. We report that LDAM are more prevalent in AD brains compared to controls and enriched in individuals with the *APOE4/4* genotype. This finding is confirmed by staining for ACSL1, a key regulator of LD biogenesis, which may serve as a useful functional protein marker of human LDAM. We also describe the induction of TG synthesis and LD accumulation by Aβ fibrils in iPSC-derived microglia (iMG), and that the LD induction is greater in isogenic APOE4/4 iMG than APOE3/3 iMG. Conditioned media from LD-high microglia induce tau phosphorylation and neurotoxicity in an APOE-dependent manner. Exposure of neurons to LD-high conditioned media increased the levels of the same TG species observed to accumulate in microglia. Since neurons in AD do not seem to upregulate *ACSL1* in AD **(Fig 1f**), we speculate that lipids accumulating in neurons are derived from microglia. This opens the possibility for a new hypothesis for LDAM-mediated pathogenesis in AD, wherein Aβ induces microglial TG lipid synthesis, lipid droplet accumulation, and subsequent secretion of neurotoxic factors in an APOE-dependent manner. In this model, these lipids might be transferred to neurons thereby inducing the hallmarks of neurodegeneration (**Fig 4i**).

While recent studies have focused on the role of APOE in astrocytes^31^, endothelial cells^32^, neurons^33^, and oligodendrocytes^34^, the LDAM expression signature, such as *ACSL1* and *NAMPT* upregulation, seems to be unique to microglia. It is likely that the APOE genotype contributes to AD pathogenesis through a number of distinct mechanisms unique to individual cell type dysfunction. A recently discovered example of this is the *APOE4-dependent* role of T-cell recruitment to the CNS and neurodegeneration in mouse tauopathy models^35^. Interestingly, this T-cell mediated neurodegeneration was not only *APOE4* dependent, but specifically dependent on microglia. We observe many chemokines secreted specifically in LD-positive microglia (**Fig. 3k),** raising the possibility that LDAM may play a role in T-cell CNS homing in AD.

A recent report showed that innate immune triggers (e.g *E.coli, Salmonella*) induce LD formation in peripheral macrophages as part of an evolutionarily ancient anti-microbial defense in which LD coated with antimicrobial proteins, such as cathelicidin (CAMP), rapidly kill bacteria^8^. We speculate that a similar program can be triggered in human microglia exposed to Aβ, LPS, and other innate immune activators, and thus impair brain homeostasis. In fact, protein aggregates found in other neurodegenerative diseases may trigger the LDAM state. For example, alpha-synuclein binding to TLR2 and TLR5 induces microglial NLRP3 inflammasome activation, which is a shared signature seen in LDAM^36^. Given that we recently identified that LDAM are abundant in the aging mouse brain, LDAMs may also be triggered by hitherto unknown protein aggregates and innate immune activators that accumulate with age. Interestingly, the most enriched pathway in human LD containing iMGs is “cellular senescence”, similar to lipid-laden “foamy macrophages” in atherosclerosis which have a senescent phenotype and are drivers of pathology^37^. Perhaps in the natural aging of various organs, LD accumulating tissue-resident macrophages represent a general class of senescent myeloid cells that are drivers of tissue inflammation.

One strategy to clear lipid droplet accumulation in microglia that we present here is PI3K inhibition, which has been shown to increase autophagy^38^. Activation of innate immune receptors such as TLR4 in microglia suppresses autophagy^39^, and this mechanism has been shown to increase LD levels in microglia^40^. Enhancing autophagy has previously been explored as a strategy to modulate neurodegeneration disease progression but with a focus on autophagic degradation of proteins^41^, as opposed to the autophagic degradation of intracellular lipid droplets (i.e. lipophagy). Future investigation into the beneficial role of specifically increasing lipophagy in AD models may better elucidate the damaging roles of lipid accumulation in AD.

In summary, we discovered that the *APOE4* genotype facilitates the microglial transition to an evolutionarily conserved, maladaptive, and damaging LDAM state in response to innate immune triggers including amyloid-beta. Future studies will have to determine whether protective *APOE* variants operate as antagonists to this microglial LDAM transition by limiting LD accumulation.

## Acknowledgments

We thank members of the Wyss-Coray lab for their support, and H. Zhang and K. Dickey for excellent laboratory management. We are grateful to the Banner Sun Health Research Institute Brain and Body Donation Program of Sun City, Arizona for the provision of human biological materials. The Brain and Body Donation Program has been supported by the National Institute of Neurological Disorders and Stroke (Beach TG; U24 NS072026 National Brain and Tissue Resource for Parkinson’s Disease and Related Disorders), the National Institute on Aging (Reiman EM; P30 AG19610 and P30AG072980, Arizona Alzheimer’s Disease Center), the Arizona Department of Health Services (contract 211002, Arizona Alzheimer’s Research Center), the Arizona Biomedical Research Commission (contracts 4001, 0011, 05-901 and 1001 to the Arizona Parkinson’s Disease Consortium) and the Michael J. Fox Foundation for Parkinson’s Research. P30 AG066515 (M.A-R. PI), and R01 AG064928-01 (Wyss-Coray PI). This work was further supported by the Knight Initiative for Brain Resilience to M.A-R. M.A-R. is Stanford Terman Fellow and a Pew-Stewart Scholar for Cancer Research, supported by The Pew Charitable Trusts and the Alexander and Margaret Stewart Trust. This work was supported by National Institute of Health P30 AG072980 (Reiman PI), R01 AG069453 (Reiman PI), P01 AG073082 (Huang PI), R03 AG071791 (Heilshorn PI, Enejder co-PI,), and R01 AG064928-01 (Wyss-Coray PI). M.S.H is supported by the National Institute on Aging (T32AG000266). R.P. is supported by the National Institute of Aging (P30AG059307). A.E. acknowledges the support of Prof. Sarah Heilshorn in the establishment of the CARS microscopy lab.

## Author Contributions

T.W.-C., M.S.H designed and conceived the experiments; G.S., T.B., E.M.R. provided human brain tissue and accompanying clinical data. M.S.H., N.S., K.S., A.S., and R.P. performed and analyzed single-nucleus RNA sequencing experiments. M.S.H., C.N.M., I.H.G., A.T., B.S., and A.S. performed human and mouse brain IF, Western blot, Oil red O, and IHC staining and imaging. N.K. provided J20 APOE KI mouse brain tissue under the supervision of Y.H. C.W. provided isogenic *APOE4/4* and *APOE3/3* iPSCs under the supervision of Y.H. M.S.H performed iMG experiments, iMG live cell imaging, and bulk RNA-seq of iMGs. C.L. and P.J. performed CARS imaging and analysis under the supervision of A.E. E.W. and J.S. performed ATAC-seq library prep and analysis under the supervision of C.G. . Y.W. prepared human iPSC-derived neurons in culture and O.Y. performed iPSC-derived neuron experiments, iPSC neuronal staining, imaging, and quantification under the supervision of Y.H. W.D. and E.R. performed lipidomics experiments under the supervision of M.A-R., C.N.M., and I.H.G. performed iMG IF and imaging. M.S.H. performed CRIPSR KO screens and analysis. M.S.H. and T.W.-C. wrote the manuscript with input from all authors. T.W.-C. supervised the study.

---

**Extended Data Figure 1.**
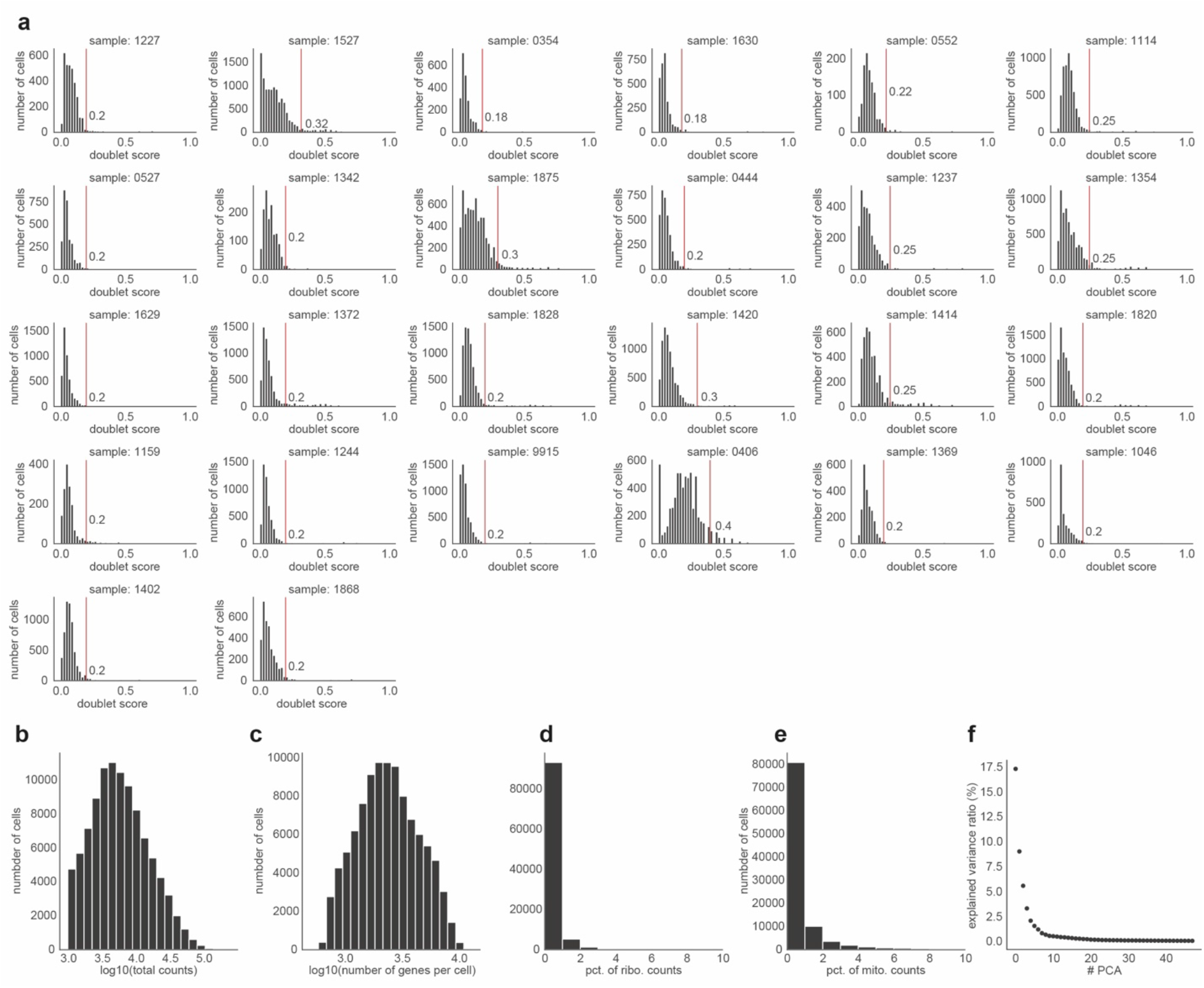
**a,** Doublet score distributions obtained with the Scrublet (0.2.3) Python package. Vertical lines indicate the doublet score thresholds identified per sample based on the distributions shown. **b-e,** Distribution of total read counts (**a**), number of genes expressed per cell (**b**), percent of reads mapped to ribosomal genes (**c**), and percent of reads mapped to mitochondrial genes (**d**) within the processed data after quality control. **f,** Percent of explained variance across the first 48 principal components.

**Extended Data Figure 2.**
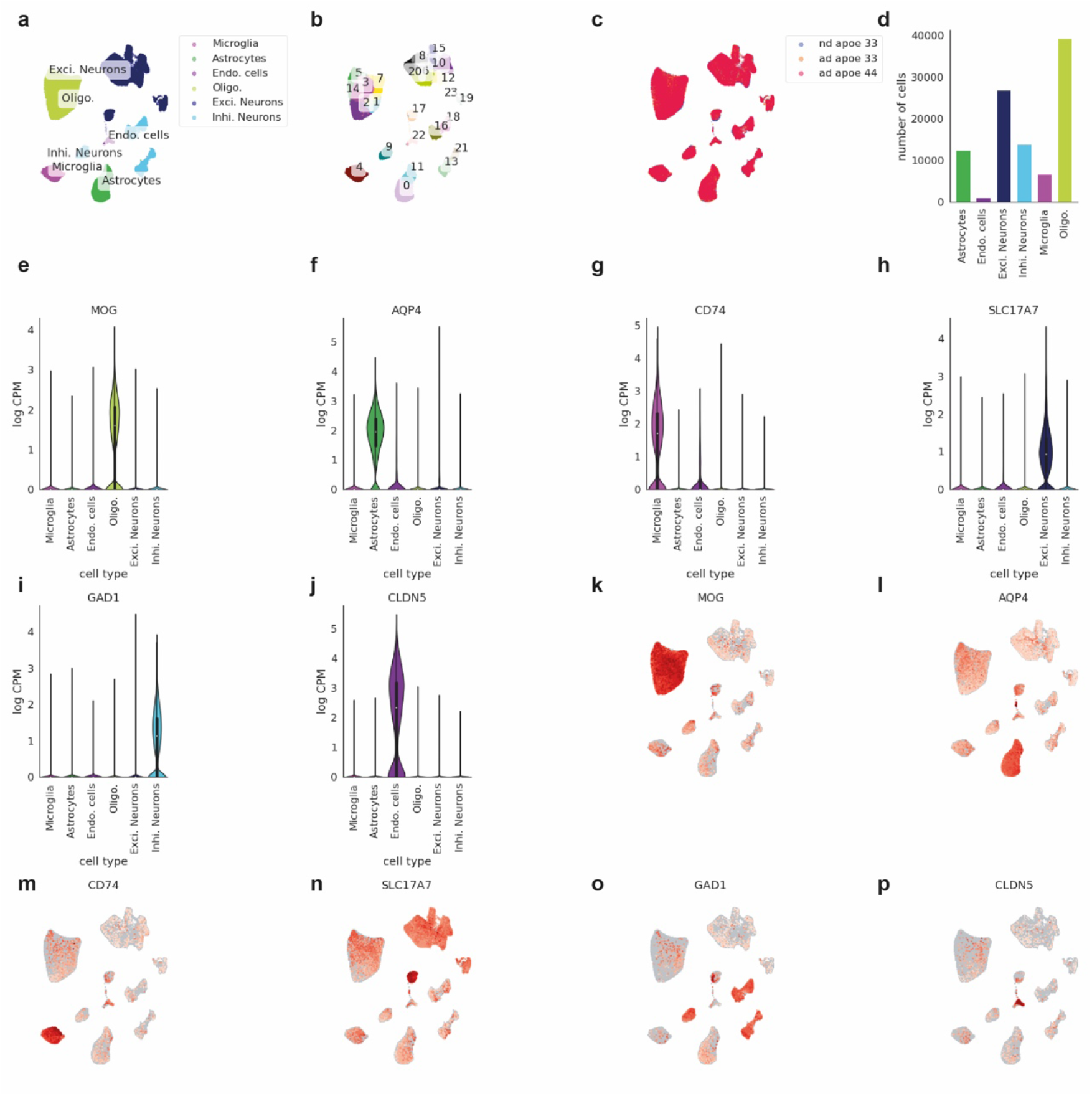
**a-c,** UMAP visualization of the whole snRNA-seq dataset after quality control and batch correction (n=100,317). Cells are colored by cell type annotation (**a**), subclusters drawn for cell type annotation (**b**), subject groups (control, AD-APOE3/3, AD-APOE4/4) (**c**). **d,** Bar chart indicating the total number of cells per cell type. **e-j,** Violin plots indicating gene expression levels of marker genes used for cell type annotations across the 6 identified cell types. **k-p,** UMAP visualization of the whole snRNA-seq dataset colored by cell type marker gene expression levels per cell.

**Extended Data Figure 3.**
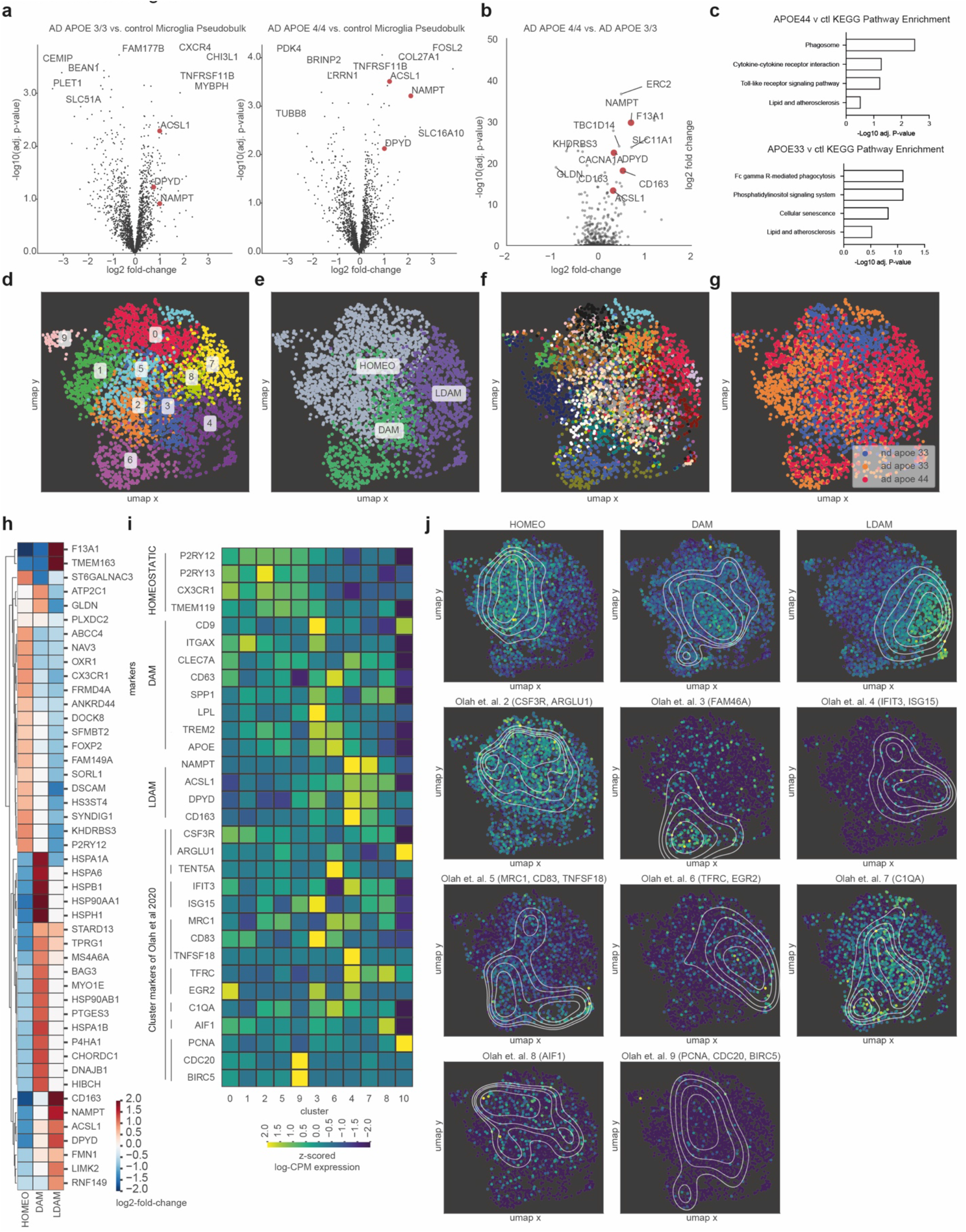
**a,** Volcano plot representing pseudobulk differential gene expression results (see Methods, Microglia pseudobulk differential gene expression) of microglia from control subjects compared to microglia (left) from subjects with AD and the APOE3/3 genotype, (right) from subjects with AD and the APOE4/4 genotype. Select lipid and metabolism-associated genes highlighted in red. **b,** Volcano plot representing single-cell differential gene expression results of microglia from subjects with AD and the APOE3/3 genotype compared to microglia from subjects with AD and the APOE4/4 genotype. Select lipid and metabolism-associated genes highlighted in red. **c**, Select KEGG pathway analysis terms and enrichment score for top 200 differential expressed genes in between control and AD-APOE4/4 microglia (top) and control and AD-APOE3/3 microglia (bottom). **d-g,** UMAP visualization of the microglia colored by identified subclusters (macrophage cluster 10 is not shown) (**d**), identified microglial states (**e**), subject IDs (**f**), and subject groups (control, AD-APOE3/3, AD-APOE4/4). **h**, Top marker genes identified for the 3 microglial states (HOMEOSTATIC, DAM, LDAM) with ‘one vs. rest’ marker identification. Single-cell differential gene expression was performed with MAST (see Methods, Single-cell differential expression). Heatmap indicates significant (adj. p-value<0.05) log2-fold changes. **i**, Normalized and z-scored gene expression levels of HOMEOSTATIC, DAM, LDAM marker genes as well as marker genes identified by Olah *et al.* (2020) across the 11 subclusters identified within the microglia. **j,** UMAP representation of microglia cells indicating signature scores per cell for HOMEOSTATIC, DAM, LDAM marker genes as well as marker genes identified by Olah *et al.* (2020). Contour lines indicate kernel density estimates of the signatures across the UMAP space.

**Extended Data Figure 4.**
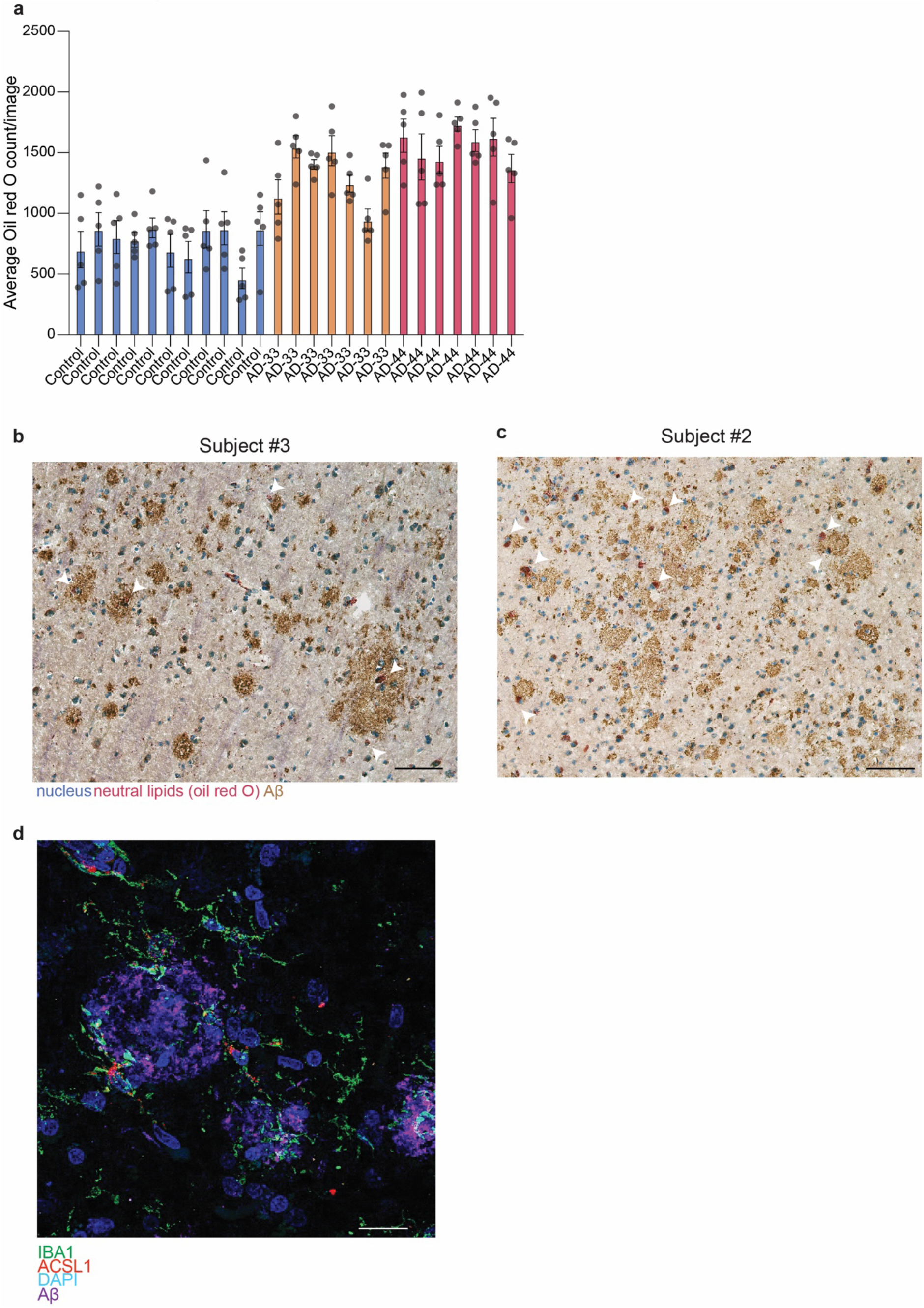
**a**, Quantification of Oil Red O counts for each subject. **b-c,** Additional Oil Red O staining of APOE4/4 AD subjects with IHC staining for Aβ. White arrowheads represent Oil Red O positive cells in or around Aβ plaques. Scale bars (black, bottom right) 50 μm. **d,** Representative immunofluorescence images of human frontal cortex adjacent to tissue used in snRNA-seq experiments stained for microglia marker IBA1 (green), ACSL1 (red), and DAPI (blue), and amyloid-beta (magenta) in an AD APOE4/4 subject. Scale bars (white, bottom right) 20 μm.

**Extended Data Figure 5.**
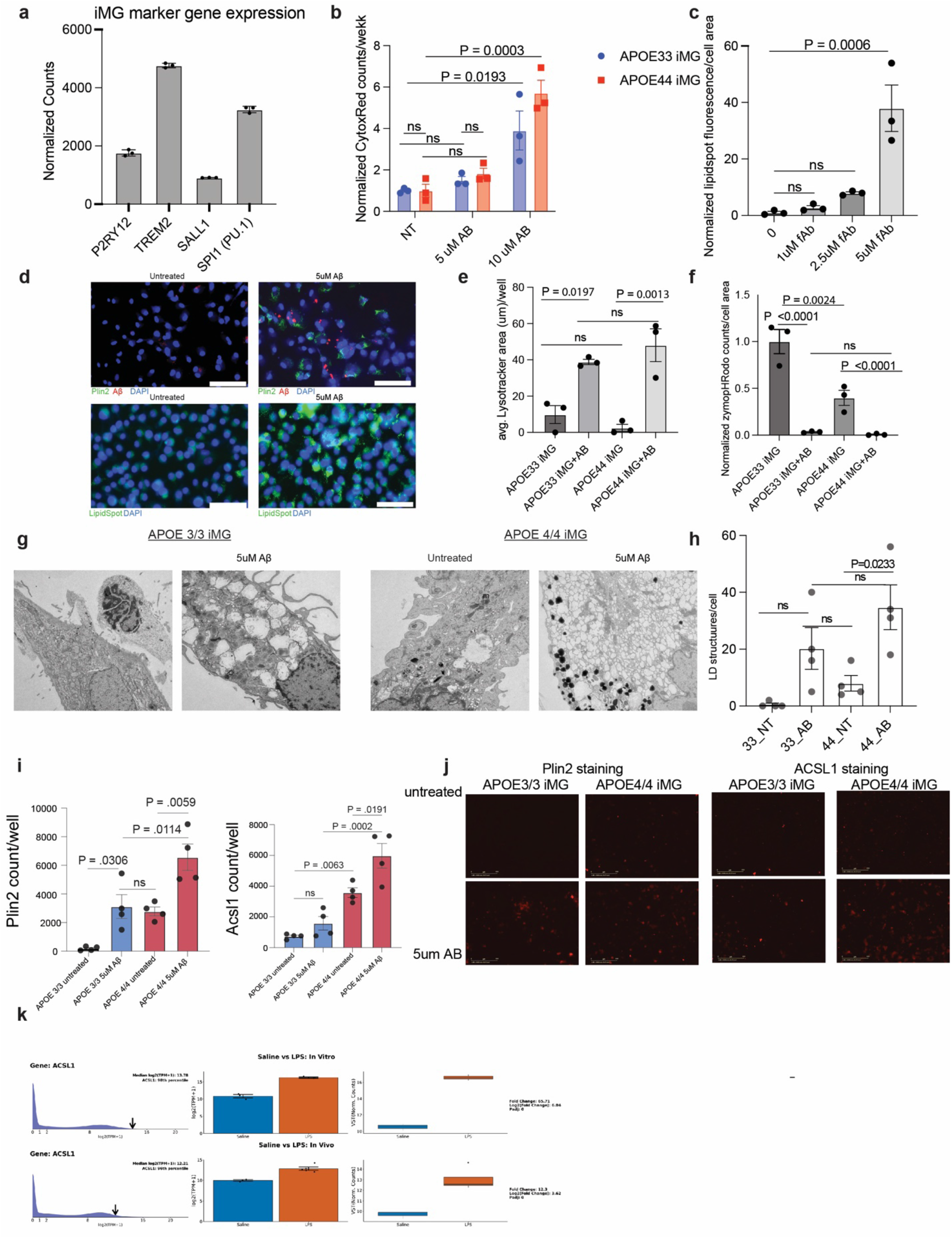
**a**, Expression of microglia marker genes in IMG. **b**, Toxicity of fAβ at different concentrations in *APOE4/4* and *APOE3/3* iMG after 24 hour incubation followed by staining with CytoxRed. Each dot represents a replicate well, error bars represent s.e.m, p-value calculated by ANOVA. **c, Dose-dependent** effect of fAβ on lipid accumulation. Each dot represents a replicate well, error bars represent s.e.m, p-value calculated by ANOVA. **d,** Representative image of human macrophages (top) untreated (left) or treated (right) with 5 μM Aβ for 24 hours (left) with PLIN2 (green) and Aβ (red) staining. Scale bars (white, bottom right) 50 um. Representative image of mouse BV2 cells (bottom) untreated (left) or treated (right) with 5uM Aβ for 24 hours (left) with LipidSpot (green) and Aβ (red) staining. Scale bars (white, bottom right) 75 μm. **e,** LysoTracker area per cell after incubation with fAβ in *APOE3/3* and *APOE4/4* iMG. Each dot represents a replicate well, error bars represent s.e.m, p-value calculated by ANOVA. **f,** Phrodo zymosan phagocytosis area per cell after incubation with fAβ in *APOE3/3* and *APOE4/4* iMG. Each dot represents a replicate well, error bars represent s.e.m, p-value calculated by ANOVA. **g-h,** Transmission electron microscopy of *APOE3/3* iMG and *APOE 4/4* iMG treated with 5 μM Aβ for 24 hours and untreated iMG. Scale bars (white, bottom right) 2 μm. Each dot represents images of individual cells, error bars represent s.e.m, p-value calculated by ANOVA. **i-j,** Representative image of *APOE4/4* iMG treated with 5 μM Aβ for 24 hours (left) with Plin2 and ACSL1 staining quantification with (**i**) with representative images (**j)** (n=4 replicate wells, error bars represent s.e.m, p-value calculated by ANOVA, Scale bars (white, bottom right) 50 μm. **k,** *ACSL1* gene expression measured in human iMG after LPS treatment as described in Hasselmann *et al*. 2019 (https://rnaseq.mind.uci.edu/blurton-jones/bulkSeq/)

**Extended Data Figure 6.**
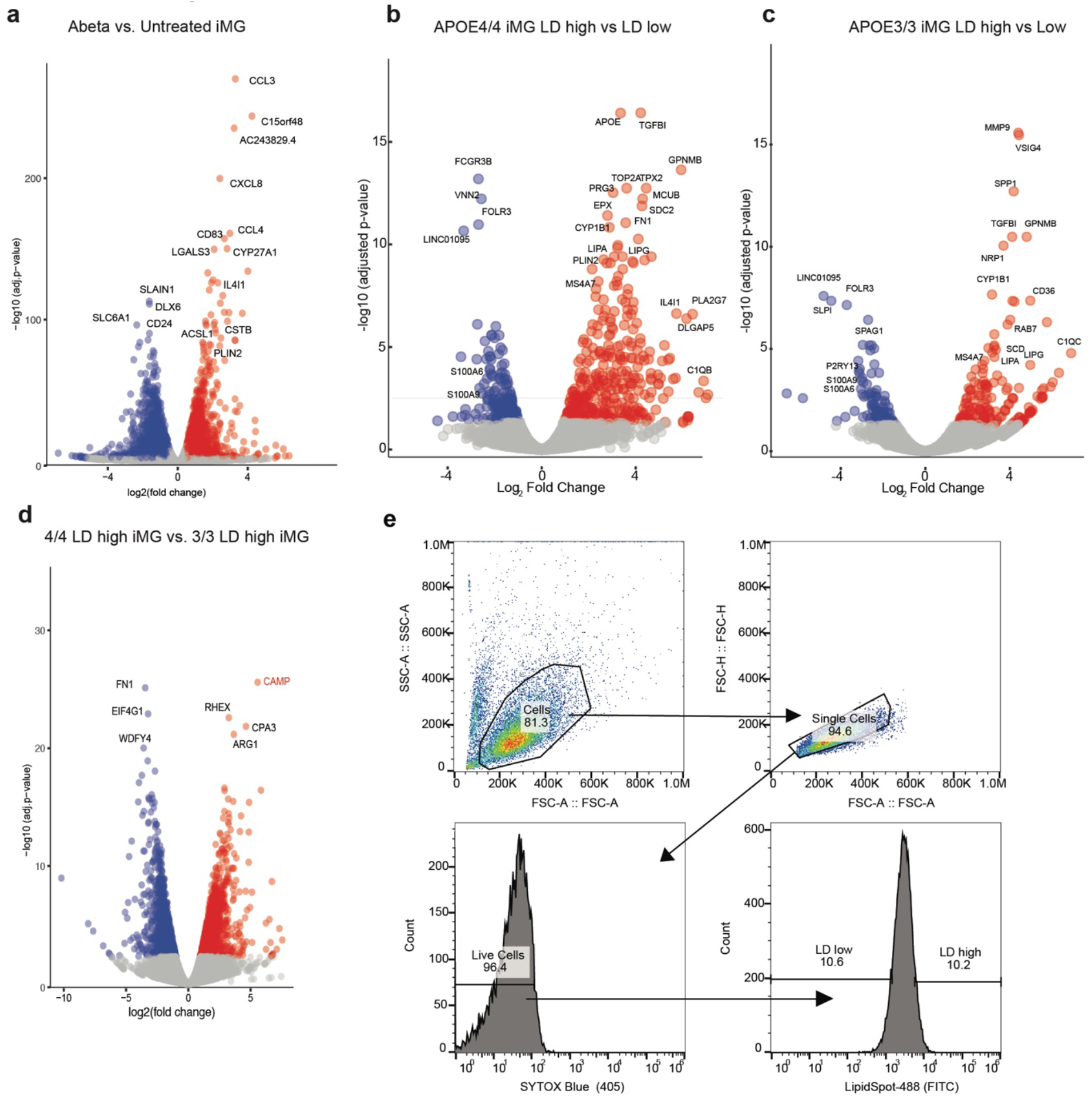
**a**, Volcano plot of differential gene expression analysis of untreated and Aβ treated *APOE4/4* iMG. **b,** Volcano plot of differential gene expression analysis of lipid droplet high and lipid droplet low *APOE4/4* iMG. **c,** Volcano plot of differential gene expression analysis of lipid droplet high and lipid droplet low *APOE3/3* iMG. **d,** Volcano plot of differential gene expression analysis of lipid droplet high *APOE3/3* iMG and lipid droplet high *APOE4/4* iMG. **e,** Example gating scheme for separating cells based on lipid droplet content.

**Extended data Figure 7.**
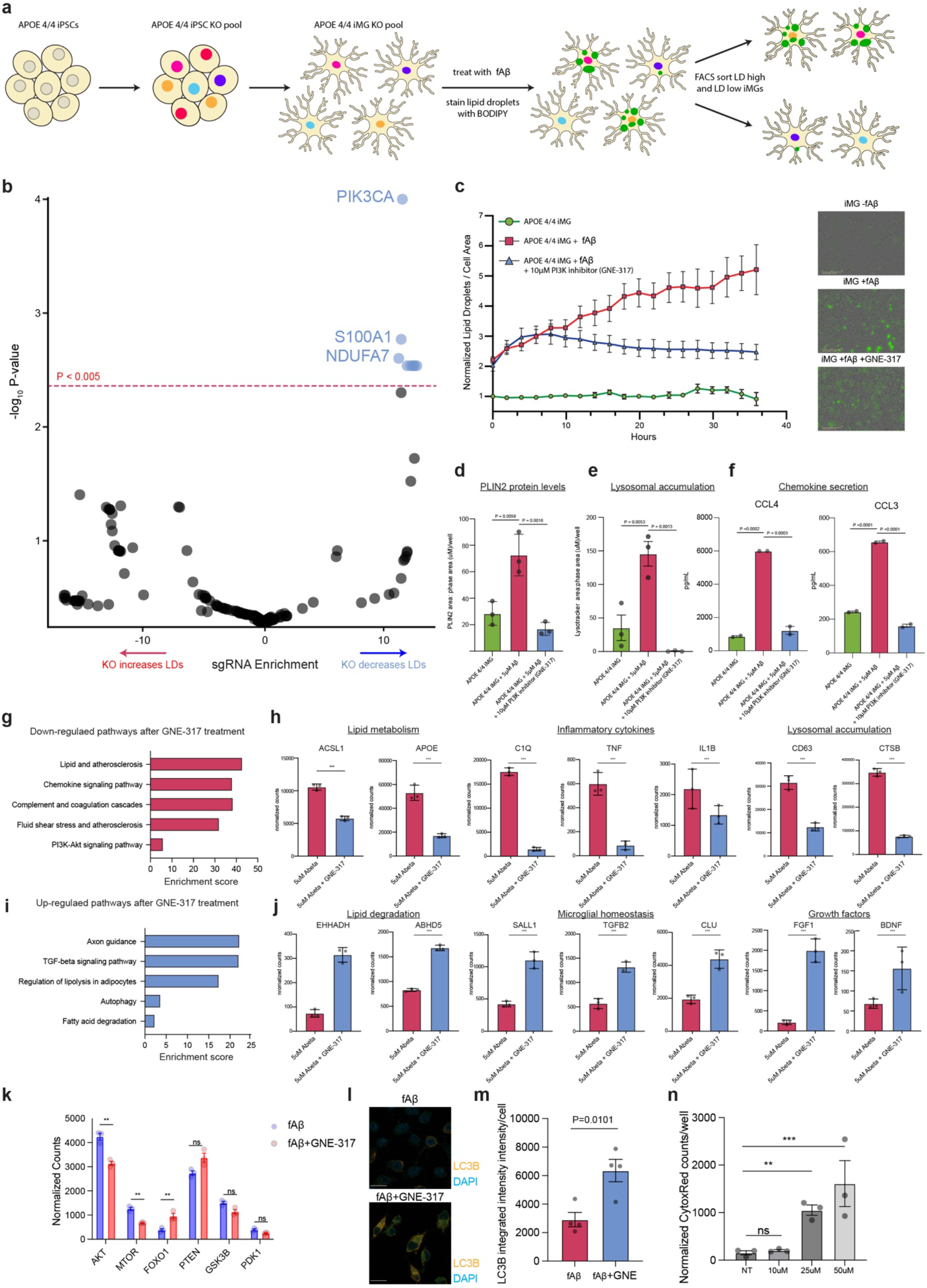
**a**, Schematic of CRISPR KO screen in *APOE4/4* iMG for lipid droplet formation following Aβ treatment. **b**, Volcano plot representing CRISPR screen results. Effect score represents log2 fold change in sgRNA counts in lipid droplet negative versus lipid droplet positive cell fraction. Screen hits with P value <0.005 colored blue. **c**, Live cell imaging of untreated *APOE4/4* iMG, 5 μM Aβ treated iMG, and 5 μM Aβ treated iMG with 10 μM GNE-317. The y axis represents average green fluorescence per cell normalized to untreated *APOE 4/4* iMG at the first time point and the x-axis represents imaging time points in hours (n=3 replicate wells per condition, error bars represent s.e.m.) (left). Representative images at the final time point (right) with LipidSpot signal represented in green. **d**, Quantification of PLIN2 staining in untreated, Aβ treated, and Aβ treated with 10 uM GNE-317 conditions iMG for 24 hours. (n=3 replicate wells, error bars represent s.e.m, p-value calculated by ANOVA). **e**, Quantification of lysotracker staining in untreated, Aβ treated, and Aβ treated with 10 μM GNE-317 conditions iMG for 24 hours. (n=3 replicate wells, error bars represent s.e.m, p-value calculated by ANOVA). **f**, Measurement of secreted chemokines in cell culture media in untreated, Aβ treated, and Aβ treated with 10 μM GNE-317 conditions iMG for 24 hours. Individual dots represent replicate wells (n=2, error bars represent s.e.m, p-value calculated by ANOVA). **g**, Select KEEG pathway enrichment terms for the top 200 significant downregulated genes ranked by p-value upon GNE-317 treatment in *APOE4/4* iMG when challenged with Aβ. **h**, Normalized gene expression counts for significantly downregulated genes with GNE-317 treatment in *APOE4/4* iMG when challenged with Aβ *APOE4/4* iMG (n=3 replicate wells, error bars represent s.e.m., *** P < 0.0001). P-values determined by DEseq2. **i**, Select KEEG pathway enrichment terms for top 200 significant upregulated genes ranked by p-value upon GNE-317 treatment in *APOE4/4* iMG when challenged with Aβ. **j**, Normalized gene expression counts for significantly upregulated genes with GNE-317 treatment in *APOE4/4* iMG when challenged with Aβ APOE4/4 iMG (n= 3 replicate wells, error bars represent s.e.m., *** P < 0.0001). P-values determined by DEseq2. **k**, Differential gene expression for genes in mTOR and autophagy pathways upon GNE-317 treatment with fAβ challenge in iMG. (n=3 replicate wells, error bars represent s.e.m., *** P < 0.0001). P-values determined by DEseq2. **l**, Representative images of LC3B staining (yellow) upon fAβ and GNE-317 treatment. The scale bar (white, bottom left) represents 20μm. **m**, Quantification of LC3B staining by integrated fluorescence intensity per DAPI signal. Individual dots represent replicate wells (n=4, error bars represent s.e.m, p-value calculated by unpaired, two-sided, t-test). **n**, Toxicity measurements of GNE-317 as determined by cytox-red. (n=4, error bars represent s.e.m, p-value calculated by ANOVA).

**Extended data Figure 8.**
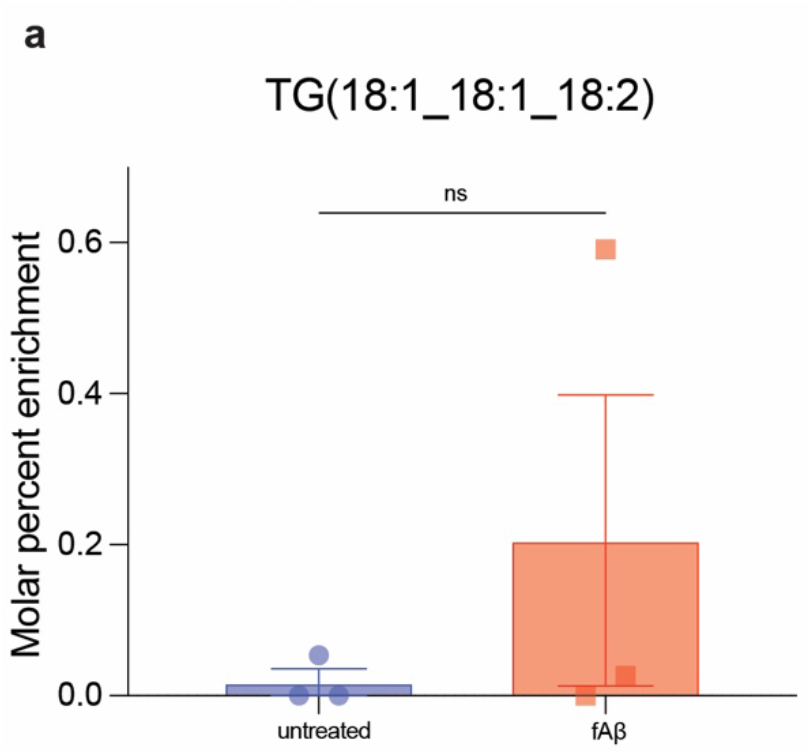
Detection of TG synthesized in microglia taken up by neurons through labeled ^13^C-glucose tracing. **a**, Measurement of ^13^C-labeled triglycerides synthesized in microglia and profiled in neurons by lipidomics after exposure to microglia conditioned media. Microglia grown in uniformly labeled ^13^C-glucose (U-13C6-glucose) were challenged with fAβ or untreated. Each dot represents an individual replicate. n=3, error bars represent s.e.m, p-value calculated by unpaired t-test.

## Methods

### Single nucleus sequencing of human brain tissue

Frozen superior frontal gyrus and fusiform gyrus tissue blocks and pathology/clinical reports were obtained from the Banner Sun Health Research Institute Brain and Body Donation Program in accordance with institutional review boards and policies at both Stanford School of Medicine and Banner Sun Health Research Institute. All samples obtained from Banner Sun Health Research Institute were stored at 80 °C until the time of processing. Isolation of nuclei from frozen brain tissue: 20-50 mg of flash frozen human brain tissue isolated from the frontal cortex was thawed in 2 mL ice-cold homogenization buffer (Molecular biology grade water, 260 mM Sucrose, 30 mM KCl, 10 mM MgCl, 20mM Tricine-KOH, 1 mM DTT, 500 μM Spermidine, 150 μM Spermine, 0.3% NP-40, Protease inhibitor, RNAse inhibitor) for 5 minutes in a pre-chilled dounce homogenizer. Tissue was dounce with “A” loose pestle for 10 strokes then dounce by 20 strokes with “B” tight pestle. The resulting homogenate was passed through a 70 micron Flowmi strainer into a prechilled 1.5mL tube. Nuclei were pelleted by centrifugation for 5 minutes at 4 °C at 350 RCF in a fixed angle centrifuge. All but 50 μL of supernatant (containing cytosolic RNA) was removed by pipetting and nuclei were resuspended in 1x homogenization buffer to a total volume of 400 μL and gently mixed by pipetting and transferred to a pre-chilled 2 mL protein-lobind tube. 400 μL of 50% Iodixanol solution (OptiPrep Sigma Catalogue # D1556) was gently mixed by pipetting with 400 μL resuspended nuclei to make a 25% iodixanol solution. 600 μL of a 30% Iodixanol solution was gently layered underneath the 25% iodixanol solution containing isolated nuclei, without mixing Iodixanol layers. Next, 600 μL of a 40% iodixanol solution was gently layered underneath the 30% iodixanol solution, without mixing iodixanol layers, resulting in a total volume of 2 mL. Samples were centrifuged in a pre-chilled swinging bucket centrifuge for 20 minutes at 4 °C at 3,000 RCF with the brake off. After centrifugation, the top 1 mL of the iodixanol layer (25% solution containing myelin and larger debris) was aspirated using a vacuum down to 1 mL total volume, containing the nuclei band. Nuclei were isolated from smaller cellular debris by removal by pipetting of the top 200 μL of the nuclei layer (at a volume between 800 μL and 1 mL in the 30%-40% iodixanol interface). 200 μL of isolated nuclei were diluted in 200 μL nuclei wash buffer (PBS with 0.1% BSA and 0.2 U/μL RNase Inhibitor. snRNA-seq libraries were prepared from nuclei using the Chromium Next GEM Single Cell 3ʹ v.3.1 according to the manufacturer’s protocol (10x Genomics). Nuclei were counted using with a TC20 Automated Cell Counter (Bio-Rad) and loaded into a 10x droplet generator at a concentration of 8,000 nuclei per sample. Thirteen pcr cycles were applied to generate cDNA, and 15 cycles for final library generation. The final snRNA-seq libraries were sequenced on a NovaSeq 6000.

### snRNA-seq Quality control

Raw gene counts were obtained by aligning reads to the hg38 genome (refdata-gex-GRCh38-2020-A) using CellRanger software (v.4.0.0) (10x Genomics). To account for unspliced nuclear transcripts, reads mapping to pre-mRNA were also counted. All subsequent analysis was implemented in Python (3.9.12) based on the Scanpy (1.9.1) single-cell data analysis package except where stated otherwise. Count data was first screened for doublets with the Scrublet (0.2.3) Python package^2^. Once each cell was doublet scored, we applied a separate doublet score threshold per sample to discard doublets from the data. Thresholds were identified between 0.15 and 0.5 per sample based on the sample-wise doublet score histograms (see Extended Figure 1a). We then applied standard filtering rules following the guideline of Luecken et al.^3^. We used the Scanpy (1.9.1) package to discard cells with (1) fewer than 500 genes or (2) less than total 1,000 reads or (3) more than 10% mitochondrial reads or (4) more than 10% ribosomal reads. Counts were then CPM scaled and log-normalized for downstream analysis.

### Global data integration and clustering

We merged then all data across the samples and used standard methods of Scanpy (1.9.1) to select the top 2,500 highly variable genes (HVGs) and calculate the top 20 principal components (PCA). The previous number of principal components were identified with the elbow method. We then used the Python implementation of BBKNN (1.5.1), a fast batch correction algorithm suitable for large datasets to integrate data across the samples^4^. BBKNN calculates a batch-corrected neighborhood graph from the imputed principal components. We set the individual sample ids as batch labels to correct for potential sample-wise batch effects. We then used the batch-corrected neighborhood graph to run Leiden clustering^5^ and to calculate a global UMAP embedding with default parameters in Scanpy (1.9.1). We used these embeddings to annotate the cells. Note that BBKNN does not modify the count data in any ways, but returns a neighborhood graph that we use for the noted downstream analyses.

### Cell type annotation

Leiden clusters were annotated one-by-one based on domain-specific expertise. We investigated each Leiden cluster separately based on the expression of common cell type markers (Exci. Neurons: *Slc17a7*, Inhi. Neurons: *Gad1*, Oligo.: *Mog*, Endothelial cells: *Cldn5*, Astrocytes: *Aqp4*, Microglia: *Cd74*) and annotated the clusters accordingly. Marker gene expression analysis was implemented in Python (3.9.12) and Scanpy (1.9.1).

### Microglia pseudobulk differential gene expression

We summed the raw counts per patient sample and hence derived ‘pseudobulk’ samples^6^ from the single-cell counts. We then used Deseq2^7^ to perform bulk data normalization and differential gene expression in R (4.3). We followed the standard Deseq2 analysis steps and conducted sequencing depth based count normalization across all APOE4/4, APOE3/3, and control samples. Then, we performed differential gene expression analysis (DGE) at standard parameters. The resulting log-scaled fold-changes were shrunken using the standard ‘apeglm’ approach. In every comparison, we discarded genes used for QC filtering (Rb* and Mt-*) from the DGE analysis since these may be biased by the quality control process. P-values were corrected with Benjamini-Hochberg procedure^8^ (FDR=0.05) per comparison.

### Microglia subsampling

In order to conduct *unbiased* single-cell level downstream analysis of the microglial cells that is not biased towards any of the patient samples, first we subsampled the microgila cluster with purification . We used the first 20 principal components calculated based on the top 2,000 highly variable genes as input and selected 1,000 cells per subject groups (control, AD-APOE3/3, AD-APOE4/4).

### Single-cell differential gene expression

We performed differential gene expression (DGE) on the subsampled data with MAST^9^ in R (4.3) by using the Seurat package. We set gender, expired age, PMI, and the percent of mitochondrial/ribosomal counts as covariates. We performed (a) pairwise comparisons (3 in total) across the three subject groups (control, AD-APOE3/3, AD-APOE4/4) (b) ‘one-vs-rest’ comparisons between the three annotated microglial states (HOMEOSTATIC, DAM, LDAM). P-values were corrected with Benjamini-Hochberg procedure^8^ (FDR=0.05) per comparison.

### Microglia subclustering

Subsampled microglia data was normalized as described previously^10–12^. Briefly, starting from the raw counts, gender, expired age, PMI, and the percent of mitochondrial/ribosomal counts were first regressed out with the *regress_out* function of the scanpy package (1.9.1). We then used the first 20 principal components based on the top 2,000 highly variable genes as input to repeat the Leiden clustering and UMAP visualization steps with default parameters in Scanpy (1.9.1) on this corrected microglia subcluster data. In order to characterize each subcluster, we calculated the mean expression level of marker genes (HOMEOSTATIC: P2RY12, P2RY13, CX3CR1, TMEM119, DAM: CD9, ITGAX, CLEC7A, CD63, SPP1, LPL, TREM2, APOE, LDAM: NAMPT, ACSL1, DPYD, CD163). We then calculated signature scores by averaging the min-max normalized expression values of these per cell. We used these signature scores to annotate the subclusters, and also mapped them to the UMAP space and performed kernel density estimation to locate them in the UMAP space (‘the single-cell landscape’).

### Prefrontal cortex immunohistochemistry and immunofluorescence

Adjacent to tissue processed for snRNA-seq was subjected to immunohistochemistry and Immunofluorescence. Prefrontal cortex of each subject were cut with a razor blade and directly submerged in 4% paraformaldehyde at 4 °C for 24 h. They were then transferred to a 30% sucrose solution in 1X PBS and stored at 4 °C until the tissue sank in its vial. Tissues were then frozen in OCT compound, stored at −80 °C, and sectioned on a cryostat (Leica CM3050S). For Oil Red O staining, slides with 10 μm sections of prefrontal cortex were removed from the −80 C freezer and brought to RT for 5 min. The slides were first incubated in propylene glycol for 5 min and then Oil Red O solution from an Abcam Oil Red O stain kit (ab150678) for 2 hr at RT. The sections were differentiated in 85% propylene glycol in distilled water for 1 min, incubated in hematoxylin for 1 min, and rinsed with multiple changes of distilled water. The slides were sealed with a glass cover slide and ProLong Gold mounting media. Imaging was performed with a ZEISS Axioskop 2 Plus microscope. Oil Red O counts per image were quantified in ImageJ and statistical analysis performed in Prism9 (Graphpad). For immunofluorescence, free-floating 50 μm sections were washed three times in PBST followed by blocking with 5% donkey serum in PBST for 1 h. Sections were incubated in PBS with 3% donkey serum and the following primary antibodies for 72 h at 4°C: goat anti-Iba1 (1:500; Abcam, ab5076), rabbit anti-ACSL1 (1:100; Thermo Fisher PA5-78713), and mouse anti-β-amyloid (1:500; Cell Signaling Technologies, #15126). After primary antibody incubation, sections were washed three times in PBST and incubated in PBS and the following secondary antibodies for 2 hours at RT: donkey anti-goat Alexa Fluor 488, donkey anti-rabbit Alexa Fluor 555, and donkey anti-mouse Alexa Fluor 647 (all 1:500; Invitrogen). Sections were incubated in DAPI (1:2000; Thermo Fisher) for 10 min, then washed three times with PBST. Sections were mounted on microscope slides with ProLong Glass mounting media. Imaging was performed with a ZEISS LSM 900 confocal microscope ZEN 3.0 (Blue Edition) software.

### Immunofluorescence staining of mouse brain sections

Human APOE3-KI and APOE4-KI mice were cross-bred with mice overexpressing mutant human APP (J20 line) to generate J20/APOE4-KI and J20/APOE3-KI mice, as we reported previously^13^. All mouse lines were maintained on a C57Bl/6J background. Sex-and age-matched wildtype mice were used as controls. Brains were collected from female J20/APOE4-KI and J20/APOE3-KI mice (n=3 for each group) at 13 months of age. Brain sections were collected (30μm) from paraformaldehyde-fixed right hemibrains on a sliding microtome fitted with a freezing stage as described previously. Free-floating 30 μm sections were washed three times in PBS followed by blocking with 10% donkey serum in PBS for 1 h. Sections were incubated in PBS with 5% donkey serum and Iba1 primary antibody (1:500; Wako 019-19741) for 72 h at 4 °C. After primary antibody incubation, sections were washed once in PBS and incubated in the following secondary antibody and lipid droplet dye for 2 hr at RT: donkey anti-rabbit 555 (1:500; Invitrogen) and LipidSpot 488 (1:1000; Biotium). Sections were incubated in DAPI (1:2000; Thermo Fisher) for 10 min, then washed three times with PBS. Sections were mounted on microscope slides with ProLong Glass mounting media. Imaging was performed with a ZEISS LSM 900 confocal microscope ZEN 3.0 (Blue Edition) software.

### iPSC maintenance and differentiation to microglia

Isogenic *APOE4/4* and *APOE3/3* iPSCs were a generated as previously described^13^. iPSCs were maintained in StemFlex medium (Gibco, A3349401) and routinely passaged as clumps onto Matrigel (Corning, 40230) coated plates. Differentiation into microglia was performed as previously described^14–16^. iPSCs were first differentiated into hematopoietic progenitor cells (HPCs) following the manufacturer’s instructions using the commercially available STEMdiff Hematopoietic Kit (Stemcell Technologies, 05310). HPCs in suspension were transferred to plates containing an adherent layer of confluent primary human astrocytes (ThermoFisher, N7805100), in media containing Iscove’s Modified Dulbecco’s Medium (ThermoFisher), fetal bovine serum (FBS) and penicillin– streptomycin and 20 ng/mL each of IL3, GM-CSF, and M-CSF (PeproTech) for 10 days. iMG were harvested and transferred into homeostatic culture conditions adapted from Muffat *et al*^16^. 30 (MGdM media) on Matrigel (Corning, 40230) coated plates for 5-15 days before assay. For assays involving staining for microscopy iMG were plated in homeostatic culture conditions (MGdM media) on Fibronectin (StemCell, 07159) coated plates to enhance cell adherence. All assays were performed under serum-free conditions (MGdM media).

### iMicroglia and macrophages Immunofluorescence and live cell microscopy

iMicroglia or human macrophages were fixed in 4% paraformaldehyde for 10 min. Cells were washed with PBS followed by blocking with 5% donkey serum in PBS for 1 h. Cells were incubated in PBS with 3% donkey serum and the following primary antibodies overnight at 4 °C: rabbit anti-Perilipin 2 (1:200; Proteintech 15294-1-AP), rabbit anti-ACSL1 (1:100; ThermoFisher PA5-78713), LC3B (1:300, ThermoFisher Scientific, PA1-46286) and mouse anti-β-amyloid (1:500; Cell Signaling Technologies, #15126). After primary antibody incubation, cells were washed with PBS and incubated in PBS and the following secondary antibodies for 2 h at RT: donkey anti-rabbit Alexa Fluor 488, donkey anti-rabbit Alexa Fluor 555, and donkey anti-mouse Alexa Fluor 647 (all 1:500; Invitrogen). Sections were incubated in DAPI (1:2000; Thermo Fisher) for 10 min, then washed three times with PBS. Cells grown and stained on coverslips were then mounted on glass microscope slides with ProLong Glass mounting media. Imaging was performed with a ZEISS LSM 900 confocal microscope ZEN 3.0 (Blue Edition) software. Additionally, for quantification of cells were fixed in replicate wells of a 96-well plate and were imaged and quantified with Incucyte S3 analysis system (Essen Bioscience). For live cell microscopy cells were untreated or treated with 5 μM Aβ fibrils, or 10 μM GNE-317 (Selleckchem, S7798) for 24 hours. After 24 hours media was changed and cells were incubated with Lipidspot 488 (biotium, 70065-T) in combination with pHrodo Red Zymosan Bioparticles (P35364, ThermoFisher) or Lysotracker (L12492, ThermoFisher), according to manufactures instructions. For Aβ fibril formation, monomeric HFIP-treated Amyloid beta protein (1-42) (Bachem cat # 4090148) was formed into fibrils as previously described^13^. Four phase-contrast, green, and red fluorescent images per well were acquired every 1 h for 36 h using an Incucyte S3 analysis system (Essen Bioscience). For each time point, red and green fluorescence was normalized to the phase confluence per well.

### Primary human macrophage cell culture conditions

Whole blood was obtained from Stanford Blood Center. Peripheral blood mononuclear cells (PBMCs) isolation was performed by diluting sample in PBS (equivalent blood volume) and transferred on top of the Ficoll layer (GE Healthcare Cat#17-5442-02). The tubes were then centrifuged at 400xg, at RT, acceleration (slow), and brake (slow) for 30 min. After centrifugation, the upper layer was discarded and the PBMCs layer at the interphase was collected in a fresh 50 mL Falcon tube. The cells were washed twice with PBS and counted. Monocytes were then isolated using the Pan Monocyte Isolation Kit (Miltenyi Biotec Cat#130-096-537) according to manufacturer instructions. Monocytes were cultured serum free X-vivo 15 media (Lonza) and differentiated into macrophages with 20 ng/mL M-CSF (Peprotech).

### BV2 cell culture conditions

Cells from the murine microglial BV2 cell line were originally obtained from Banca Biologica e Cell Factory, IRCCS Azienda Ospedaliera Universitaria San Martino, Genua, Italy. Cells were maintained in DMEM (Life Technologies) supplemented with 5% FBS and antibiotics (penicillin 100 U ml−1, streptomycin 100 U ml−1 (pen/strep) (Gibco), 10mM glutamax (Gibco) under standard culture conditions (95% relative humidity with 5% CO2 at 37 °C). Adherent cells were split using 1× TrypLE (Gibco). Experiments with deuterated lipids were conducted by replacing glucose with 10mM D-glucose (U-13C6, Cambridge isotope laboratories CLM-1396-PK) for 24h hours prior to treatment with 5 μM Aβ fibrils or no treatment.

### Primary rat microglia cell culture conditions

Primary rat microglia were cultured as previously described^14^. P10-20 rats were transcardially perfused with ice-cold PBS and immediately after perfusion, brains were rapidly dissected and placed into ice-cold PBS. Brain material was minced and transferred to an ice-cold dounce homogenizer (Wheaton) with ice-cold PBS containing 200 μL of 0.4% DNaseI per 50 mL of PBS. Tissue chunks were subjected to three successive rounds of 3 to 10 gentle strokes of the homogenizer piston and centrifuged. PercollPlus was added during centrifugation to separate myelin from cells. The cell suspension volume was adjusted to 33.4 mL with PBS and 10 mL of 100% isotonic Percoll (9 mL Percoll PLUS (GE Healthcare), 1 mL 10x PBS without Ca and Mg, 9 μL 1 M CaCl2, 5 μL 1 M MgCl2) was added and thoroughly mixed (23% isotonic Percoll final). Suspensions were centrifuged (15 min, 500g, 4°C) and the supernatant and top layer of myelin were discarded. Rat CD11b/c (Microglia) MicroBeads (Miltenyi, cat. no. 130-105-643) were used to enrich rat microglia using MACS columns according to manufacturers’ instructions. Serum-free rat microglia basal microglial growth medium was sterile-filtered and stored at 4°C for up to one month and was comprised of: DMEM/F12 containing 100 units/mL penicillin, 100 μg/mL streptomycin, 2mM glutamine, 5 μg/ml N-acetyl cysteine, 5 μg/ml insulin, 100 μg/mL apo-transferrin, and 100 ng/mL sodium selenite, ovine wool cholesterol (1.5 μg/mL, Avanti Polar Lipids), heparan sulfate (1 μg/mL, Galen Laboratory Supplies). The final medium was comprised of basal media containing human TGF-β2 (2 ng/mL, Peprotech), murine IL-34 (100 ng/mL, R&D Systems),). Cells were plated on 24-well plates coated with poly-D-lysine (Gibco). Cells were grown in a humidified incubator held at 37°C and 10% CO2. 50% medium changes were performed every 2 to 3 days. For live cell microscopy cells were untreated or treated with 5 μM Aβ fibrils (as described above) pre-incubated with an Aβ antibody (Cell Signaling Technologies, #15126) conjugated to an Alexa Fluor 555 secondary (Invitorogen), along with Lipidspot 488 (Biotium, 70065-T) and imaged for 24 hours.

### Preparation of Slides for iPSC-derived Cultures for CARS imaging

The cell cultures, adherent to circular cover glasses, were positioned on secondary #1.5H cover glasses (Thorlabs) with a thin layer of phosphate-buffered saline (PBS). To circumvent interference with lipid CARS signal generation, neither mounting media, antifade agents, nor surfactants were employed. The edges of the cover glass were sealed with VALAP, a homogeneous blend of equal parts petroleum jelly, lanolin, and paraffin, which served to prevent sample desiccation.

### Characterization by CARS and Confocal Fluorescence Microscopy

Intracellular lipids in iPSC-derived cells were visualized and quantified using coherent anti-Stokes Raman scattering (CARS) microscopy. An inverted microscope (Nikon, Ti2-E equipped with a C2 confocal scanning head and a Nikon CFI Apochromat TIRF 100XC oil immersion objective) was utilized for this purpose. The C2 scanner was retrofitted with a slidable mirror (Optique Peter), allowing for a convenient switch between fluorescence excitation using the laser diodes (at wavelengths 405, 488, 561, and 647 nm) and CARS excitation. In CARS imaging mode, carbon–hydrogen (C–H) vibrations were coherently driven by temporally and spatially overlapping two near-infrared laser beams, generated by a picosecond-pulsed laser system (APE America Inc., picoEmerald S with 2 ps pulse length, 80 MHz repetition rate, and 10 cm^−1^ bandwidth) consisting of a 1031 nm mode-locked ytterbium fiber laser and an optical parametric oscillator (OPO) tunable between 700-960 nm (pumped by the second harmonic of the 1031 nm laser). The OPO wavelength was set to 797 nm to drive the symmetric stretching vibration of CH_2_ at 2850 cm^−1^. The quadratic dependence of the CARS signal on the number density of the probed C–H vibrational group rendered sharp contrast for the lipid-dense regions without requiring external labels or disruptive sample preparations. The CARS signal generated by simultaneously scanning the two excitation beams over the sample was detected pixel-by-pixel with a photomultiplier tube (Hamamatsu, R6357) in the forward direction with optical filters that minimized background signals (Semrock; two FF01-640-20, one FF01-750/SP). The excitation powers at the sample position were 18 mW for the pump (OPO) beam and 15 mW for the Stokes (1031 nm) beam. For the iPSC-derived microglia-like cells, 5 to 16 image stacks were acquired for each APOE genotype and amyloid beta treatment condition. Each stack comprised 19 slices at a resolution of 1024 × 1024 pixels (77.14 × 77.14 μm) per image with a dwell time of 10.8 μs/pixel. Slices were spaced 0.4 μm apart, yielding a total imaging depth of 7.2 μm. Cell-specific immunohistochemistry facilitated the identification of the microglia (IBA1) with confocal fluorescence, which was collected in the same positions as the nonlinear imaging with a dwell time of 5.3 μs/pixel. The microglia were stained with Alexa Fluor 488. For selected APOE/treatment combinations, a CARS spectrum was acquired in the C– H stretching region for one field of view by varying the OPO wavelength between 785.5 and 801 nm, with 0.5 nm intervals per acquisition, amounting to a total of 32 image stacks making up the spectrum. All images were analyzed using the Fiji distribution package of ImageJ.^[1]^ The CARS stacks underwent east shadows correction, Gaussian blur filtering (σ=1), and background subtraction (30 pixel rolling ball radius). Lipid particles were identified by thresholding using the FindFoci method, with a search parameter for half peak value of 0.1, peak parameter for minimum peak height relative to background of 0, and a minimum peak size of 200 pixels.^[2]^ Cells were identified by thresholding with the Triangle method.^[3]^ Intracellular lipid features were quantified using the “3D Objects Counter” command. To generate spectral line plots, the spectra from all individual intracellular lipid features in each cell were normalized and averaged.

### iMicroglia secreted Cytokine/Chemokine assay

APOE4/4 iMG were treated with 5 μM Aβ fibrils stained with Lipidspot 488 as described above then top 10% highest and 10% lowest Lipidspot fluorescent cells were seeded into 96-well plates at 5,000 cells per well by FACS (Sony, MA900) in 100 μl MGdM media for 24 h. Media supernatant was collected, and secreted signaling proteins were measured in culture supernatants by the Human Immune Monitoring Center (HIMC) at Stanford University using a Human 48-plex Luminex Procarta Immuno-assay (ThermoFisher).

### ATAC-seq library preparation

Approximately 100,000 iMG were pelleted at 300 xg and 4 °C. Next, pellets were gently resuspended in ice-cold 50 μL lysis buffer (10 mM Tris-HCl pH 7.4, 10 mM NaCl, 3 mM MgCl2, 0.1% IGEPAL CA-630) and spun down at 500 xg for 10 min and 4 °C. The supernatants were discarded, and pellets gently resuspended in 50 μL of transposition reaction mix (25 μL tagment DNA buffer (Nextera, Illumina), 2.5 μL tagment DNA enzyme (Nextera, Illumina), 22.5 μL nuclease free water) and incubated at 37 °C for 30 minutes. Tagmented DNA was purified using MinElute PCR purification kit (Qiagen) and size selected for 70 – 500 bp using AmpureXP beads (Beckman Coulter). Libraries were constructed and amplified using 1.25 μM Nextera index primers and NEBNext High-Fidelity 2xPCR Master Mix (New England BioLabs). A quantitative PCR was run to determine the optimal number of cycles. Libraries were gel size selected for 165-300 bp fragments and single-end sequenced for 100 cycles (PE100) on an Illumina NovaSeq6000.

### ATAC-seq analysis

Bowtie2 with default parameters was used to map ATAC-seq. HOMER was used to convert aligned reads into ‘‘tag directories’’ for further analysis^15^. ATAC-seq experiments were performed in replicate and peaks were called with parameters -L 0 -C 0 -fdr 0.9 -minDist 200 - size 200. IDR was used to test for reproducibility between replicates. IDR peaks were merged using HOMER’s mergePeaks and annotated with HOMER’s annotatePeaks.pl with a size parameter of 250. Differential enhancer peaks (+/- 3kb from TSS) were identified using DESeq2 with FC>1 and p-adj <0.05. HOMER’s motif analysis (findMotifsGenome.pl) including known default motifs and de novo motifs was used to identify motifs enriched in enhancer peak regions over background. The background sequences were from random GC matched genome sequences. The UCSC genome browser was used to visualize ATAC-seq data.

### iMicroglia bulk RNA-sequencing

For Aβ treated conditions, iMG were exposed to 5 μM Aβ fibrils for 24 hours or DMSO before RNA isolation. For LD high vs low conditions iMG were exposed to 5 μM Aβ fibrils for 24 hours, stained with LipidSpot 488 as described above and top 10% highest and 10% lowest Lipidspot fluorescent cells were separated by FACS (Sony, MA900). iMicroglia RNA was isolated from the cell pellets using an RNeasy Plus Micro kit (Qiagen, 74034). mRNA was transcribed into full-length complementary DNA using a SMART-Seq v4 Ultra Low Input RNA kit (Clontech) according to the manufacturer’s instructions. Full-length cDNA (150 pg) was processed using a Nextera XT DNA library preparation kit (Illumina) according to the manufacturer’s protocol. Library quality was verified using the Agilent 2100 Bioanalyzer and the Agilent High Sensitivity DNA kit. Sequencing was carried out using an Illumina NexSeq 550, paired-end, 2× 100 bp depth sequencer. Reads were mapped to the human hg38 reference genome using STAR (v.2.5.1b). Raw read counts were generated with STAR using the GeneCounts function. Differential expression in RNA-seq was analyzed using the package R DESeq2.

### Electron microscopy

Cells were grown on aclar and then fixed in Karnovsky’s fixative: 2% Glutaraldehyde (EMS Cat# 16000) and 4% para-formaldehyde (EMS Cat# 15700) in 0.1M Sodium Cacodylate (EMS Cat# 12300) pH 7.4 for 1 hr, chilled and sent to Stanford’s CSIF on ice. They were then post-fixed in cold 1% Osmium tetroxide (EMS Cat# 19100) in water and allowed to warm for 2 hr in a hood, washed 3X with ultra-filtered water, then bloc stained 2 hours in 1% Uranyl Acetate at RT. Samples were then dehydrated in a series of ethanol washes for 10 minutes each @ RT beginning at 30%, 50%, 70%, 95%, changed to 100% 2X, then Propylene Oxide (PO) for 10 min. Samples are infiltrated with EMbed-812 resin (EMS Cat#14120) mixed 1:1, and 2:1 with PO for 2 hrs each. The samples are then placed into EMbed-812 for 2 hours opened then placed into flat molds w/labels and fresh resin and placed into 65 °C oven overnight. Sections were taken around 90 nm, picked up on formvar/Carbon coated Cu grids, stained for 40 seconds in 3.5% Uranyl Acetate in 50% Acetone followed by staining in Sato’s Lead for 2 minutes. Observed in the JEOL JEM-1400 120kV and photos were taken using a Gatan Orius 2k X 4k digital camera.

### GW lipid droplet CRISPR–Cas9 screen

U937 cells were acquired from the American Type Culture Collection (CRL-1593.2). Cells were maintained in suspension culture using spinner flasks for library propagation and tissue culture plates for single-gene knockout lines, all in sterile-filtered U937 growth medium (RPMI-1640 supplemented with 2 mM glutamine, 100 units mL–1 penicillin, 100 mg mL–1 streptomycin, and 10% heat-inactivated FBS. Cells were cultured in a humidified 37 °C incubator set at 5% CO2. A 10-sgRNA-per-gene CRISPR/Cas9 deletion library (Human CRISPR Knockout library was a gift from Michael Bassik (Addgene # 101926-101934)) was infected into Cas9-expressing U937 cells as described^16^. Briefly ∼300 million U937 cells stably expressing SFFV-Cas9-BFP were infected with the ten-guide-per-gene genome-wide sgRNA library at a multiplicity of infection <1. Infected cells underwent puromycin selection (1 μg mL–1) for 5 days, after which puromycin was removed and cells were resuspended in normal growth medium without puromycin. After selection, sgRNA infection was measured as >90% of cells as indicated by measuring mCherry-positive cells with flow cytometry. For the lipid droplet screen, the 10% FBS in the media was replaced with 10% aged (>80 years of age) human plasma pooled from multiple donors, dialyzed, and delipidated to better represent the aged circulating monocyte environment. U937s will then stained with BODIPY 493/503 (1:2,000 from a 1 mg/mL stock solution in DMSO; Thermo Fisher) and top 10% highest and 10% lowest BODIPY fluorescent cells were separated by FACS (Sony, MA900). Screens were performed in duplicate. At the end of each screen genomic DNA was extracted for all screen populations separately according to the protocol included with QIAGEN Blood Maxi Kit. Using known universal sequences present in the lentivirally incorporated DNA, sgRNA sequences were amplified and prepared for sequencing by two sequential PCR reactions as described^16^. Products were sequenced using an Illumina Nextseq to monitor library composition (30–40 million reads per library). Trimmed sequences were aligned to libraries using Bowtie, with zero mismatches tolerated and all alignments from multimapped reads included. Guide composition and comparisons across bound and unbound fractions were analyzed using castle^16^ version 1.0. Enrichment of individual sgRNAs was then calculated as a median-normalized log ratio of the fraction of counts, as described^16^. For each gene, a maximum likelihood estimator was used to identify the most likely effect size and associated log-likelihood ratio (confidence score) by comparing the distribution of gene-targeting guides to a background of non-targeting and safe-targeting guides.

### iMG CRISPR–Cas9 screen

iMG CRISPR-Cas9 screens were performed using modified single guide RNA (sgRNA) lentiviral infection with Cas9 protein electroporation (SLICE) approach^43^. The human CRISPR Knockout library was a gift from Michael Bassik (Addgene #101927). In brief, ∼40 million *APOE4/4* iPSCs were lentiviral infected with the 10 guide/gene sgRNA sub-libraries at a multiplicity of infection <1. Infected cells underwent puromycin selection (.8 μg/mL) for 2 days after which point puromycin was removed and cells were resuspended in normal growth media without puromycin. iPSCs were differentiated into iMG as described above. At day 15 of iMG-human astrocyte co-culture, iMG cells were pelleted and resuspended in Lonza electroporation buffer P3 (Lonza, V4XP-3032) at 5 million cells/100mL. Cas9 protein (MacroLab, Berkeley, 40mM stock) was added to the cell suspension at a 1:10 v/v ratio. Cells were electroporated at 5 million cells/100 mL cells per cuvette using a Lonza 4-D nucleofector with pulsecode DP-148 (Lonza, VVPA-1002). Cells were co-cultured for 5 additional days with human astrocyte before transferring cells to homeostatic culture conditions (MGdM media) as described above. In duplicate culture conditions cells were treated with 5 uM Aβ fibrils for 24 hours, stained with BODIPY 493/503 (1:2,000 from a 1 mg/mL stock solution in DMSO; Thermo Fisher) and top 10% highest and 10% lowest BODIPY fluorescent cells were separated by FACS (Sony, MA900). Genomic DNA was extracted for all populations separately using a QIAGEN Blood Midi Kit, sgRNA sequences were amplified by PCR using common flanking primers, and indices and adaptors were attached to amplicons in a second PCR. Deep sequencing of sgRNA sequences on an Illumina Nextseq550 was used to monitor library composition. Guide composition was analyzed and compared to the plasmid library and between conditions using casTLE (https://bitbucket.org/dmorgens/castle). In brief, casTLE compares each set of gene-targeting guides to the negative controls, comprising non-targeting and non-genic (‘safe-targeting’) sgRNAs, which have been shown to more aptly control for on-target toxicity owing to endonuclease-induced DNA damage. The enrichment of individual guides was calculated as the log ratio between LD high and LD low populations, and gene-level effects were calculated from ten guides targeting each gene. P values were then calculated by permutating the targeting guides as previously described^17^.

### Neuronal differentiation of hiPSCs

hiPSCs were derived into neurons as previously described^16^, with slight modifications to increase yield. hiPSCs were dissociated with Accutase followed by quenching with warm (37°C) N2B27 medium. N2B27 medium consisted of 1:1 DMEM/F12 (11330032, Thermo Fisher) and Neurobasal Media (21103049, Thermo Fisher), 1% N2 Supplement (21103049, Thermo Fisher), 1% B27 (17504044, Thermo Fisher), 1% MEM Non-essential Amino Acids (11140050, Thermo Fisher), 1% Glutamax (35050061, Thermo Fisher), and 0.5% penicillin–streptomycin (15140122, Thermo Fisher). Dissociated hiPSCs were then pelleted and resuspended in embryoid body media (10µM SB431542 (1614, Tocris) and 0.25µM LDN (04-0074, Stemgent) in N2B27) with 10µM ROCK inhibitor (1254, Tocris), followed by growth in suspension in a T-75 flask (12-565-349, Fisher Scientific). For the first 3 hours, the flasks were shaken manually once per hour. On days 2, 4, and 6, the media was replaced with fresh embryoid body medium without ROCK inhibitor. On day 8, spheres were plated as neural progenitors onto a 10cm dish precoated with growth factor reduced (GFR) Matrigel (CB-40230A, Fisher Scientific). Neural progenitors were allowed to form neuronal rosettes and sustained in N2B27 media alone for days 8–15. During this period, half of the media was replaced every 48–72 hours depending on confluency and media consumption. On Day 16, the neuronal rosettes were lifted using STEMdiff™ Neural Rosette Selection Reagent (05832, StemCell Tech) and plated onto 3 wells of a 6-well plate precoated with GFR Matrigel in N2B27 with 100ng/ml FGFb (100-18B, Peprotech) and 100ng/ml EGF (AF-100-15, Peptrotech). This N2B27 medium with FGFb and EGF was replaced daily. On day 20, the neural progenitors were passaged by dissociating with Accutase, quenching with N2B27, and resuspending in STEMdiff™ Neural Progenitor Medium (05833, StemCell Tech) at 1.2×10^6^ cells/2ml for 1 well of 6-well plate, precoated with GFR Matrigel. For days 21-27, neural progenitor cells were fed with fresh Neural Progenitor Medium every day. On day 28, the neural progenitor cells were dissociated with Accutase, N2B27 media was added to bring the volume of cell suspension to 40ml, and cells were filtered through a 40µm cell strainer (08-771-1, Fisher). Cells were then collected by centrifugation and resuspended in complete neuronal medium. Neuronal media consisted of 10ng/ml BDNF (450-02, Peprotech) and 10ng/ml GDNF (450-10, Peprotech) in N2B27 with 10nM DAPT (2634, Tocris). Next, the cells were counted and plated at a concentration of 2×10^5^ cells/well onto 12mm coated glass coverslips (354087, Corning) in a 24-well plate. Coverslips were coated with poly-L-lysine (P4707, Sigma-Aldrich) and mouse-Laminin (23017015, Gibco) prior to plating. 50% of culture medium was replaced on maturing neurons every 3–4 days. DAPT was removed after the first week. Experiments were performed on neuronal cultures that had been differentiated 4 weeks.

### Lipid droplet accumulating microglia (LDAM) media treatment of neurons

LDAM-high and LDAM-low conditioned media from 1 million APOEE4/4, APOE3/3, and APOE-KO iMG were prepared by exposing iMG to 5 μM fAβ for 24-hours, washed 3x with PBS, stained with Lipidspot 488 as described above then top 10% highest and 10% lowest Lipidspot fluorescent cells were sorted by FACS (Sony, MA900) and grown in N2B27 medium (as described above) supplemented with 100 ng/ml IL-34, 10 ng/ml CSF1 and 10ng/ml TGF-beta for 12 hours. Conditioned media was collected by centrifuging media supernatant at 2,000g for 10 min to remove cell debris. 10% of the total volume from the iMG conditioned media was added to fresh N2B27 with 10ng/ml BDNF (450-02, Peprotech) and 10ng/ml GDNF (450-10, Peprotech) prior to treating neurons. To treat neurons with the various LDAM media, N2B27 neuronal media was completely removed from the cells and replaced with the iMG conditioned media containing GDNF and BDNF. After 48 hours, the neurons were fixed for immunocytochemistry.

### Immunocytochemistry, imaging, and quantification

Neurons were washed with 1X DPBS (14080055, Gibco) and fixed in 4% paraformaldehyde for 15 minutes. The neurons were then washed 3 times for 5 minutes with 1X DPBS (14080055, Gibco) containing 0.1% Tween. Next, the neurons were permeabilized and blocked with a 1-hour wash in 1X DPBS (14080055, Gibco) containing 10% Normal Donkey Serum (017000121, Jackson Immuno) and 0.5% Triton-X. Cells were then stained with primary antibodies overnight at 4°C targeting the following proteins: MAP2 (PA1-10005, ThermoFisher Scientific, 1:5000), AT8 (MN1020, ThermoFisher Scientific, 1:500), and Caspase-3 (9661, Cell Signaling Technology, 1:500). The secondary antibodies were IgG conjugated to Alexa Fluor 488 (donkey anti-mouse, A-21202, Life Technologies Corporation, 1:1000), Alexa Fluor 594 (donkey anti-rabbit, A-21207, Life Technologies Corporation, 1:1000), and Alexa Fluor 647 (donkey anti-chicken, 703-605-155, Jackson Immuno, 1:500). Coverslips were mounted to microscope slides with VECTASHIELD Prolong Gold with DAPI (H-1200-10, Vector Labs). Images were taken with a FV3000 confocal laser scanning microscope (Olympus) at 20X or 40X. Image analysis to quantify AT8 or Caspase-3 positivity in hiPSC-derived neuron stains was performed using custom macros written in the open-source Fiji (ImageJ) software. For all image analyses, a standard threshold value was chosen and automatically applied to each channel of each image prior to measurement. For quantification of AT8 staining, the total area of AT8 staining was normalized to the total area of MAP2 staining. For quantification of Caspase-3 staining, the total count of Caspase-3 positivity was normalized to the total count of DAPI positivity.

### Preparation of mouse cortical primary neurons

Newborn mouse pups on the first day after birth were humanely euthanized through decapitation, and their cortices were carefully collected using microdissection techniques. The freshly obtained cortices were then rinsed with a dissection medium (Thermo Fisher Scientific, 14170161) before being subjected to tissue dissociation using scissors, 0.25% trypsin, and pipette trituration. Cell strainers with a pore size of 70 µm were utilized to remove any remaining tissue fragments from the digested cortices. The dissociated neurons were then plated onto 24-well plates with coverslips coated with poly-l-lysine (Newcomer Supply, 1339A) using minimum essential medium (Thermo Fisher Scientific, 21010046) supplemented with 10% inactivated fetal calf serum (Thermo Fisher Scientific, 10438026), 2 mM glutamine, and penicillin and streptomycin (Thermo Fisher Scientific, SV30010). After 24 hours of initial seeding, a complete medium change was performed using neurobasal medium (Thermo Fisher Scientific, 21103049) supplemented with B-27 (Invitrogen, 17504044) and 2 mM GlutaMax (Thermo Fisher Scientific, 35050061). The primary neuronal cells were maintained at a temperature of 37 °C with a 5% CO2 environment. The detailed protocol can be found via 10.1038/s41586-022-05221-y.

### Lipid extraction and lipidomics

Dried lipids were reconstituted by a buffer consisting of 50 μL of ACN:IPA:water in a ratio of 13:6:1 (v/v/v). The mixture was then vigorously mixed using a vortex for a duration of 10 minutes at a temperature of 4 °C. Afterward, the samples were centrifuged at maximum speed for 10 minutes at 4 °C. A volume of 45 μL of the supernatant was then carefully transferred into glass insert vials for further analysis using LC/MS.

Lipid profiling was conducted using an ID-X tribrid mass spectrometer equipped with a heated electrospray ionization (HESI) probe. C18-based lipid separation was performed using an Ascentis Express C18 column coupled with a guard column. The mobile phases consisted of ammonium formate and formic acid dissolved in water, acetonitrile, and 2-propanol. The mass spectrometer parameters, including temperatures, resolutions, voltages, and gas settings, were optimized for lipid analysis. HCD fragmentation and data-dependent tandem mass spectrometry (ddMS2) were used for comprehensive lipid identification. LipidSearch and Compound Discoverer software were utilized for unbiased differential analysis and lipid annotation. Metabolite abundance was accurately quantified using TraceFinder (Thermo Fisher Scientific). Mass tolerance of 5 ppm was applied for the extraction of ion chromatograms, ensuring accurate measurement of lipid levels. The full detail of the method can be found via 10.1073/pnas.2219953120.

## Data availability

sn-RNAseq data, iMG RNA-seq data, and iMG CRISPR screening data will be made available through interactive web-app and raw sequencing data will be publicly deposited to GEO (Gene Expression Omnibus) upon publication.

## Code availability

Data analyses and graphing have been carried out using free available software packages. All code associated with the single-nucSeq data analyses will be released as reproducible Jupyter Notebooks upon publication.

